# Olivar: automated variant aware primer design for multiplex tiled amplicon sequencing of pathogens

**DOI:** 10.1101/2023.02.11.528155

**Authors:** Michael X. Wang, Esther G. Lou, Nicolae Sapoval, Eddie Kim, Prashant Kalvapalle, Bryce Kille, R. A. Leo Elworth, Yunxi Liu, Yilei Fu, Lauren B. Stadler, Todd J. Treangen

## Abstract

Tiled amplicon sequencing has served as an essential tool for tracking the spread and evolution of pathogens. Over 2 million complete SARS-CoV-2 genomes are now publicly available, most sequenced and assembled via tiled amplicon sequencing. While computational tools for tiled amplicon design exist, they require downstream manual optimization both computationally and experimentally, which is slow and costly. Here we present Olivar, a first step towards a fully automated, variant-aware design of tiled amplicons for pathogen genomes. Olivar converts each nucleotide of the target genome into a numeric risk score, capturing undesired sequence features that should be avoided. In a direct comparison with PrimalScheme, we show that Olivar has fewer SNPs overlapping with primers and predicted PCR byproducts. We also compared Olivar head-to-head with ARTIC v4.1, the most widely used primer set for SARS-CoV-2 sequencing, and show Olivar yields similar read mapping rates (∼90%) and better coverage to the manually designed ARTIC v4.1 amplicons. We also evaluated Olivar on real wastewater samples and found that Olivar had up to 3-fold higher mapping rates while retaining similar coverage. In summary, Olivar automates and accelerates the generation of tiled amplicons, even in situations of high mutation frequency and/or density. Olivar is available as a web application at https://olivar.rice.edu. Olivar can also be installed locally as a command line tool with Bioconda. Source code, installation guide and usage are available at https://github.com/treangenlab/Olivar.

## Introduction

The devastating COVID-19 pandemic has forever highlighted the utility and importance of biosurveillance for tracking the spread of emerging pathogens. Metagenomic sequencing of environmental samples has enabled the discovery of novel pathogens^1^, provided real-time insights into the spread and evolution of infectious disease^2^, and enabled the exploration of variant-specific effects on the host^3^. However, the relatively high cost and long turnaround time of metagenomic sequencing remain impractical when a large number of samples and fast turnaround times are necessary, which is the baseline for monitoring pathogens from the environment. Targeted approaches offer advantages in this setting as the ratio of the targeted pathogen is typically low compared to non-target background sequences^4,5^ (e.g., sequencing of wastewater samples by^6^), making untargeted metagenomic sequencing both more expensive, computationally intensive, and often lacking in sensitivity.

Thus, targeted amplification or enrichment can both decrease sequencing cost and improve the sequencing sensitivity for pathogen genomes of interest^4,5^. PCR tiling and DNA hybridization probes are two common approaches to amplify whole genomes of specific viruses^5,7^. While DNA hybridization probes are better at preserving the relative abundance of different species, PCR tiling has the advantage of simpler experimental workflow, faster turnaround time, and less DNA input requirement^7^. Therefore, PCR tiling has been widely used to monitor SARS-CoV-2 and characterize SARS-CoV-2 variant^8^. As of December 27th, 2022, there are 14.4 million SARS-CoV-2 genomes on GISAID^9^, and 6.5 million available in NCBI GenBank^10^, the vast majority of which were sequenced and assembled via tiled amplicon sequencing. However, when combining hundreds of primers within a single tube, PCR tiling has similar pitfalls as multiplexed PCR, including (i) uneven amplification of different genomic regions and (ii) excessive PCR byproducts (e.g., primer dimers and amplification of non-targeted sequences), resulting in a higher cost to reach a minimum acceptable sequencing depth^11^. PCR byproducts can also result in false variant calls^12^, requiring manual oversight, re-analysis and slow down the deployment which is critical in the midst of the pandemic. Moreover, PCR primers should be designed to avoid genomic regions with heavy variation^13^ (e.g., single-nucleotide polymorphisms or SNPs) and secondary structures^11^ to prevent amplicon dropout. Altogether, these pitfalls can lead to higher sequencing cost, uneven coverage, and lower sensitivity, which to date requires one or more runs of experimental validation and manual primer redesign, making the development of a PCR tiling assay costly and labor intensive^14^.

Although there are existing tools for designing PCR tiling, some design tiled amplicons as single plex assays^15^ or a number of small primer pools (<10 primers)^16^, instead of multiplexed assays where tens or hundreds of primers are mixed in the same reaction. Furthermore, previous approaches do not optimize all of the aforementioned criteria simultaneously, nor do they adequately explore the solution space of possible primer combinations^4^. The current state of the art PCR tiling design software tool, PrimalScheme^4^, takes a sequential approach to primer design. Specifically, starting from the left side (5’ end) of the genome, PrimalScheme sequentially designs each primer until the whole genome is covered, thus, newly designed primers will not affect the choice of previously designed primers. Although PrimalScheme also considers genomic variations, GC content, primer dimers, etc., the choice of new primers will be limited by previously designed primers. For example, the region where a new primer is generated might have high GC content, or candidates of the new primer might form primer dimers with previously designed primers. In the worst-case scenario for PrimalScheme, a gap in tiling will exist since no primer candidate satisfies the design requirements, leading to reduced genomic coverage and/or requiring manual redesign. Thus, their output primers are semi-optimized and often require further tweaking and redesign^14^. This is evidenced by the most widely used PCR tiling primer set in ARTIC^17^, initially designed with PrimalScheme and used to sequence millions of SARS-CoV-2 genomes at this point. ARTIC has undergone several iterations of manual tweaking and optimization^13,14,17^, including the primer dimer issue of the latest ARTIC v4.1^12^, and will continue to require manual tweaking and refinement as new variants arise.

Here we present Olivar, an end-to-end pipeline for rapid and automatic design of primers for PCR tiling. Olivar accomplishes this by introducing the concept of the “risk” of primer design at the single nucleotide level, enabling fast evaluation of thousands of potential tiled amplicon sets. Olivar looks for designs that avoid regions with high-risk scores based on SNPs, non-specificity, GC contents, and sequence complexity. We selected these four components according to known challenges with multiplexed primer design: SNPs represent sequence variation, non-specificity represents the likelihood of non-specific priming, while sequences with extreme GC content and/or low complexity are more likely to be repetitive, bearing secondary structures^18^ and producing more primer dimers^11^. Olivar also implements the SADDLE algorithm^11^ to optimize primer dimers in parallel and provides a separate validation module that allows users to evaluate their multiplex PCR primers from various aspects, including the likelihood of dimerization, amplification of non-targeted sequences, etc.

To evaluate the performance of our method, we used Olivar to automatically design a set of primers that tile the entire SARS-CoV-2 genome in 146 amplicons in under 30 minutes. In a direct in-silico comparison with PrimalScheme, Olivar had lower predicted primer dimerization, fewer SNPs overlapping with primers (4 vs. 18), and fewer predicted non-specific amplifications (5 vs. 27). We conducted an experimental head-to-head comparison with the latest ARTIC v4.1, the most widely used tiled amplicons for SARS-CoV-2, and found that Olivar had similar mapping rates (∼90%) and better coverage for synthetic RNA samples with both low (18) and high (35) cycle threshold (Ct) values.

We also tested Olivar and ARTIC v4.1 on 4 wastewater samples, and showed that Olivar has 1 to 3-fold higher mapping rates and similar coverage. Furthermore, Olivar includes an interactive visualization module that shows the targeted sequence’s risk landscape and the primer placement, allowing a convenient overview of the automated design (Figure 1).

**Figure 1.**
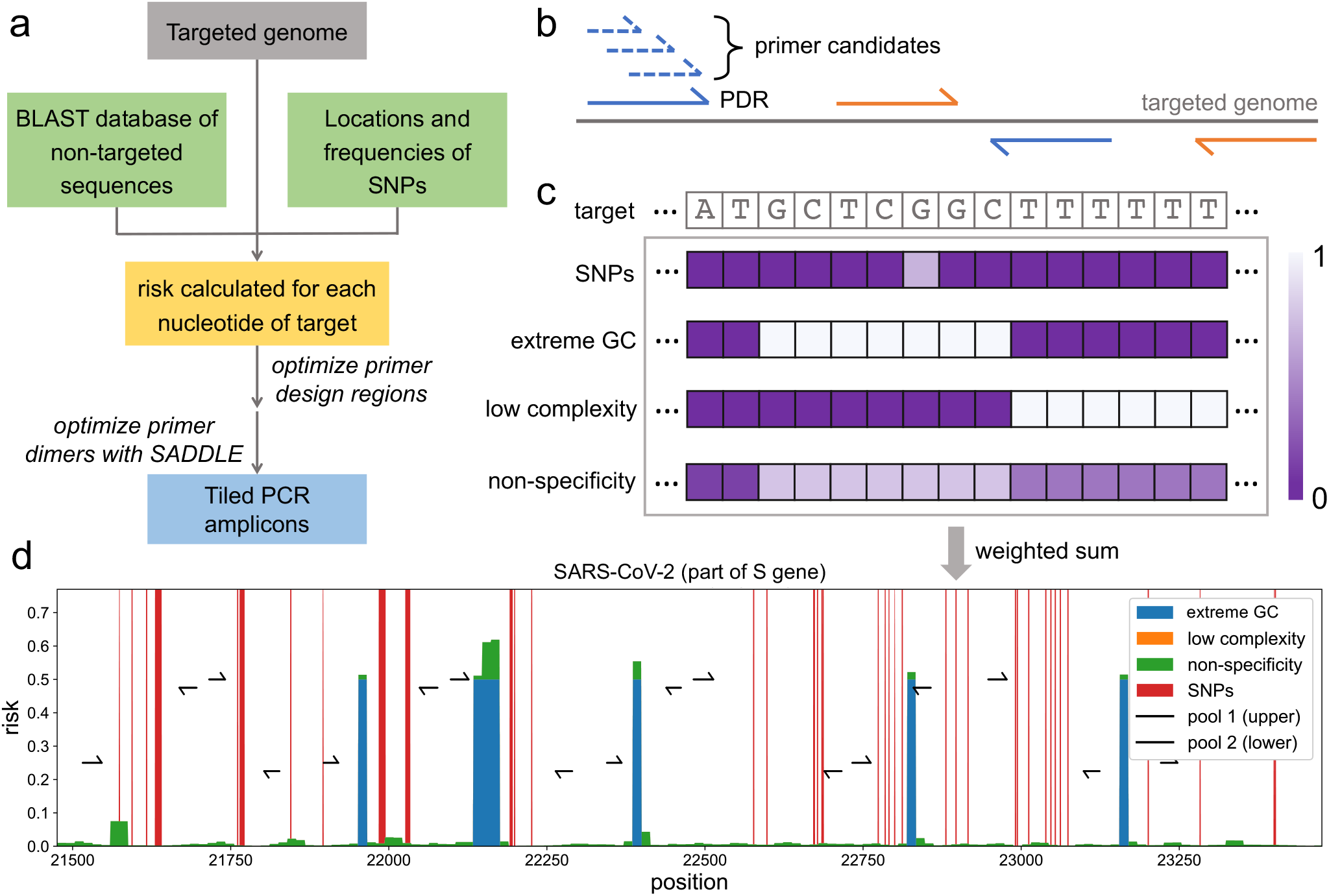
Overall workflow of Olivar and example output. **(a)** The input of Olivar consists of three components: the targeted sequence, a BLAST database built with non-targeted sequences and single nucleotide polymorphisms (SNPs) to be avoided by primers. Based on the provided inputs, a risk score is calculated for each nucleotide of the targeted sequence. Primer design regions are optimized according to the array of risk scores. One or more primer candidates are generated for each primer design region, and the primer set with minimum primer dimerization is selected with the previously published algorithm SADDLE. **(b)** A primer design region (PDR) is a short region (40nt by default) on the targeted genome. A pair of PDRs (blue or orange solid lines) covers the genomic region between them, and a valid set of PDRs should cover the whole targeted genome. Primer candidates (blue dashed lines) are generated from each PDR by SADDLE. PDRs are assigned into two pools (pool 1 in blue and pool 2 in orange) to avoid overlapping amplicons. **(c)** A risk array consists of four components: SNPs, extreme GC content, low sequence complexity and non-specificity, and all of them are calculated for each nucleotide of target and range between 0 and 1. The final risk is calculated as the weighted sum of the four risk components. **(d)** An example of the risk landscape of the SARS-CoV-2 genome, as well as primers designed by Olivar. The beginning of the S gene is shown, and each risk component is shown in a different color. Different risk components shown in the figure are stacked together instead of overlapping. Primers are assigned into two pools to avoid overlapping amplicons. Together the two pools cover the whole genome.

## Results

There are three major steps in Olivar, as illustrated in Figure 1a,

1. Generation of a risk score for each nucleotide of the targeted sequence based on user-provided inputs (reference sequence, location of SNPs, BLAST database, etc.).
2. Generation and evaluation of primer design region (PDR) sets based on a Loss function.
3. Generation of primer candidates for each PDR and optimization of primer dimer with the SADDLE algorithm.

The primer design region (PDR) is a short DNA sequence (40nt by default) from which primer candidates are generated with SADDLE (Figure 1b). The risk score emphasizes sequence features that should be avoided for primer design (e.g., high GC regions), and its definition can be tailored for specific applications. Here, we select four components critical to primer design to calculate the risk score: single-nucleotide polymorphisms (SNPs), high/low GC content (extreme GC), homopolymers (low complexity), and repeated sequence across genomes (non-specificity). Each nucleotide is given a score for each of the four sequence features, creating four different arrays named as risk components. Risk components are then weighted with user-defined weights and summed together to generate the risk array (Figure 1c). The risk array can be visualized as a risk landscape and overlaid with the primers designed, giving users a better understanding of how primers are placed on the targeted sequence (Figure 1d). The visualization module is included in the Olivar software as well as the Olivar web app. Given an array of risk scores, Olivar can efficiently evaluate thousands of potential PDR sets and choose the best one based on a Loss function. The best PDR set is input to SADDLE and the combination of primer candidates with minimum dimerization likelihood is output as the final primer set. Detailed description of risk components, generation of PDR sets and the Loss function can be found in Methods.

### Optimization of PDRs

The location of PDRs is crucial for primer design since there is little room for optimizing primer candidates if a PDR overlaps with an undesirable sequence feature (e.g., high GC region). The risk array serves as a guidance for the placement of PDRs, enabling rapid and quantitative evaluation of a set of PDRs by a customized Loss function. The above framework is compatible with various applications involving nucleotide sequence design, and it is not restricted to PCR tiling. However, PCR tiling could take the most advantage of the risk array since there is no explicit requirement for the location of PDRs as long as the targeted sequence is fully covered, leaving plenty of space for optimization. On the contrary, applications such as mutation detection with real-time PCR (qPCR) may require primers to be designed around a certain position of the genome, leaving fewer space for PDR optimization. Other qPCR applications, for example pathogen detection, might not have such restrictions. For a set of PDRs that is “valid” for PCR tiling (here, valid is defined as targeted sequence is fully covered with inserts, where an insert is the sequence between a pair of PDRs), the risk of each PDR as well as the Loss of the PDR set can be readily calculated, allowing thousands of PDR sets to be evaluated within an acceptable amount of time. Specifically, risk of a PDR is the sum of risk scores within it, while the Loss of a PDR set is determined by the risk of the worst PDRs. The definition of the Loss function is based on our experience that the performance of a multiplexed PCR assay is usually undermined by a few “bad players”, or primers bearing undesired features such as extreme GC content, low sequence complexity, etc.^11^ Instead of calculating the total risk of all PDRs, we focus the Loss function on the worst PDRs to prevent such “bad players”. Detailed description about risk of PDRs and the Loss function can be found in the Method section. Intuitively, one could randomly generate a large number of PDR sets and choose the one with the lowest Loss. To accelerate the optimization process, we introduced a certain level of greediness into the random generation, with details described in Methods. In short, given that PDRs should not overlap with each other, as well as the desired range of amplicon length, a PDR must fall into a certain region of the targeted sequence (Figure 2a). Instead of randomly selecting a PDR within that region, high risk PDRs are excluded from the random selection, with high risk defined as PDR risk greater than *X* th percentile. We experimented *X* from 10 to 90, and as *X* becomes greater, more greediness is introduced (Figure 2bc). Here the reference genome for Sars-Cov-2 is used (GISAID^9^ ISL_ID: EPI_ISL_402124), with other input data and parameters described in Methods. While the results indicate that smaller *X* is better, there is a higher chance of overfitting. Hence, we set *X* as 30 for better universality. Detailed description of generating a valid PDR set can be found in the Method section.

**Figure 2.**
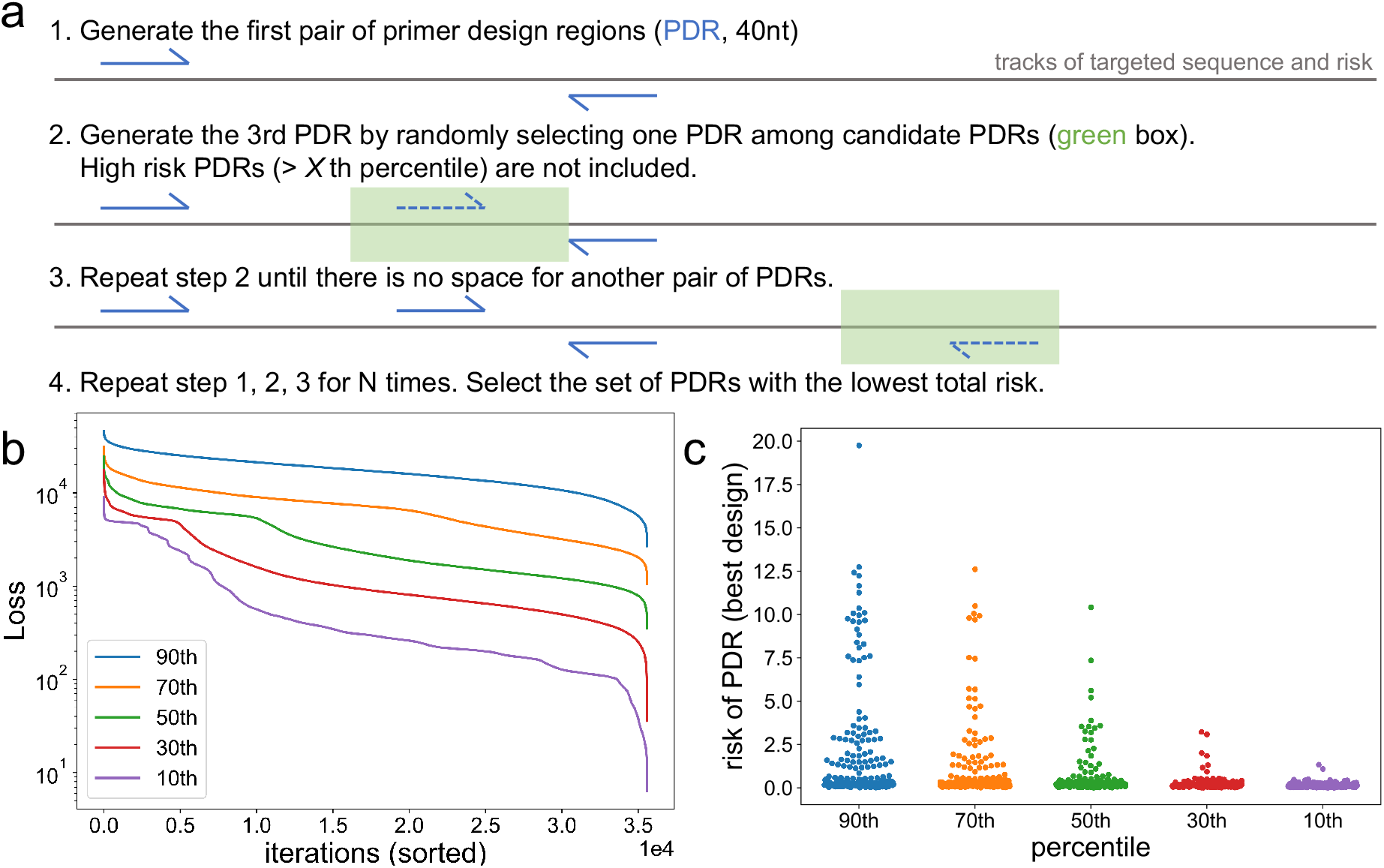
Optimization of primer design regions (PDRs). **(a)** Starting from the 5’ end of the targeted sequence, a pair of PDRs is randomly generated. The length of each PDR is fixed to 40nt. Given that PDRs should not overlap with each other, as well as the desired range of amplicon length, a PDR must fall into a certain region of the targeted sequence, as indicated by the green box. The risk of each 40-mer within this region is calculated as the sum of all single nucleotide risk. Low risk 40-mers (≤*X* th percentile) are considered as candidate PDRs and the next PDR is randomly selected among them. PDRs are repeatedly generated until there is no space for another pair of PDRs, and the total risk of the PDR set is calculated. The whole process is repeated N times, with N determined by desired amplicon length and the length of targeted sequence, and the PDR set with the lowest total risk is selected for downstream design. **(b**,**c)** Tuning the threshold for determining candidate PDRs, with Sars-Cov-2 genome as the targeted sequence. Detailed description of other design parameters can be found in the methods section. *X* is set as 30 for following designs. **(b)** For each percentile threshold, 35,584 PDR sets are generated, and their Loss is sorted. **(c)** For each percentile threshold, the PDR set with the lowest Loss is selected and the risk of each PDR is shown.

### Olivar designs PDRs that effectively avoid highly variable genomic regions

To demonstrate Olivar’s ability to avoid high-risk regions in a target genome, we used the publicly available SARS-CoV-2 data on Nextstrain^19^ where Shannon’s entropy of each base in the reference genome is provided (Figure 3a). Calculation of Nextstrain entropy can be found in Methods. We input locations with entropy greater than 0.01 to Olivar as an analogy of SNP location and frequency and optimized PDRs with default parameters described in Methods. Compared to a naively generated set of PDRs, the Olivar optimized PDR set has about 10-fold lower Loss (765.4 vs. 7644.0, Figure 3b), after 35,584 iterations on a personal computer (2.4GHz 8-Core CPU) in 20 minutes wall clock, with peak memory usage less than 500MB. Figure 3c shows the histogram of entropy of all bases in the reference genome, as well as bases within naive PDRs or Olivar PDRs, and every base with entropy greater than 0.3 is avoided by Olivar.

**Figure 3.**
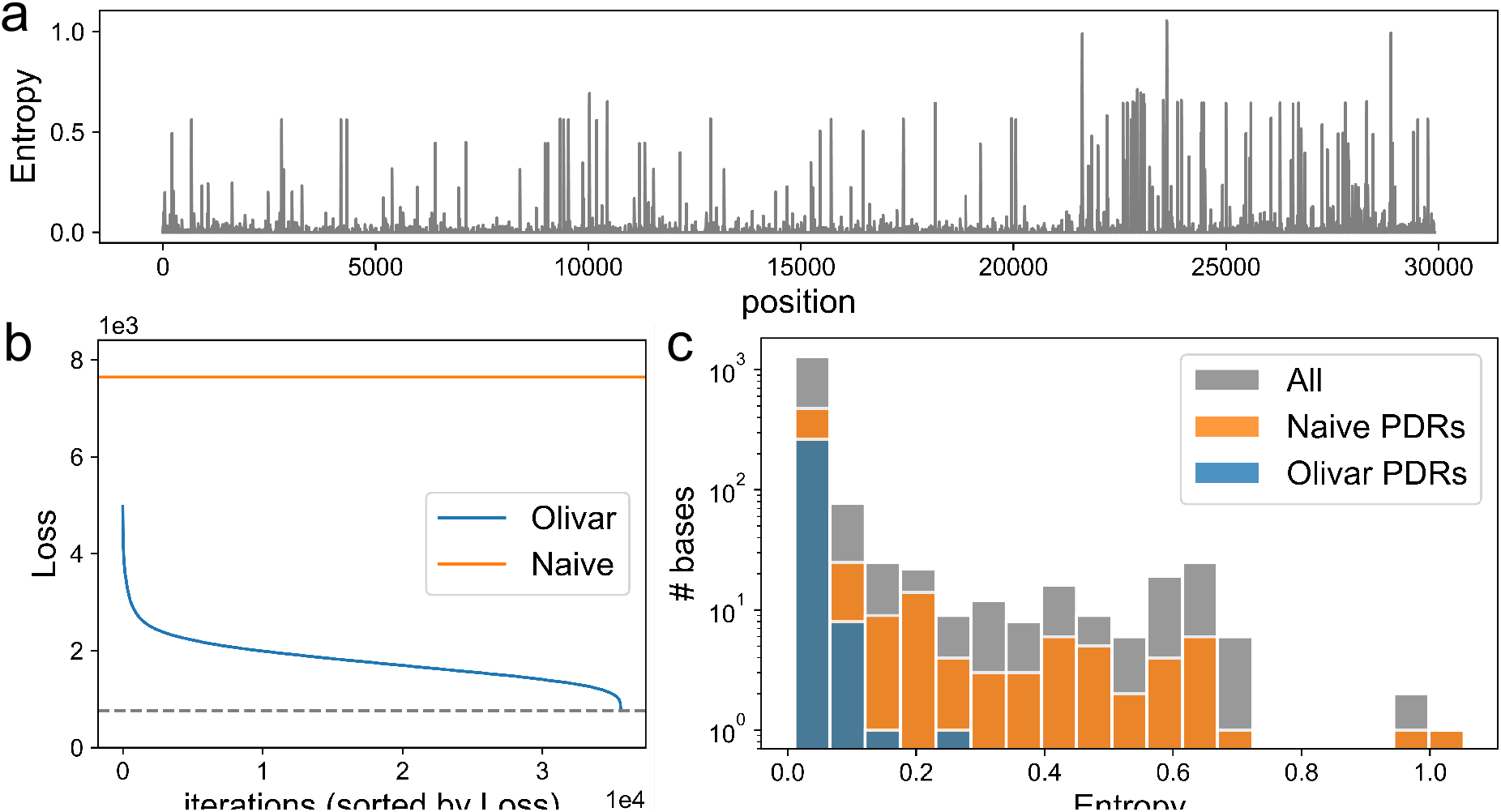
Optimization of PDRs with nucleotide entropy provided by the Nextstrain SARS-CoV-2 database. **(a)** Shannon entropy at each base position, provided by Nextstrain. Details about entropy calculation can be found in Methods. Entropy larger than 0.01 (1,517 bases) is input to Olivar as SNP frequency. **(b)** 35,584 PDR sets are generated by Olivar and the optimal PDR set with minimal Loss of 765.4 is chosen (dashed line). A randomly generated naive set of PDRs has a Loss of 7644.0. **(c)** Histogram of entropy of bases. Of the 1,517 bases with entropy greater than 0.01 (gray), 275 overlap with Olivar PDRs (blue), while 560 overlap with naive PDRs (orange).

### Olivar outperforms PrimalScheme in silico

We used Olivar and the state-of-the-art PCR tiling design software PrimalScheme to design tiled amplicons for SARS-CoV-2, based on public genomes on GISAID before March 1st, 2022. We first generated SNPs of Delta (B.1.617.2) and Omicron (B.1.1.529) variants with the software tool Variant Database^20^. 440 SNPs with frequencies greater than 1% were used as input to Olivar and PrimalScheme. Since PrimalScheme takes a group of sequences (no more than 100 sequences) as input instead of locations and frequencies of SNPs, we created 100 pseudo genomes bearing those 440 SNPs as input to PrimalScheme. Human genome assembly GRCh38.p13 was used as the non-targeted BLAST database for Olivar. Desired amplicon length was set to 252nt to 420nt for Olivar and PrimalScheme, with other input parameters kept as default. Since there is randomness in the Olivar pipeline, specifically in optimizing PDRs, we conducted 6 runs to test its reproducibility. On the other hand, there is no randomness in the PrimalScheme pipeline and PrimalScheme will generate the same primer set with the same input. We first compared 1 of the 6 Olivar designs with the Primalscheme design (Figure 4a-d). Compared with PrimalScheme, Olivar primers had a lower predicted dimerization likelihood, represented by the dimer_score of SADDLE^11^(Figure 4a,b), as well as lower BLAST hits against human genome (Figure 4c). The risk score of each primer was also calculated, with Olivar having a lower average risk per primer (0.14 vs. 0.28, Figure 4d). Across 6 Olivar runs, Olivar primers had fewer SNPs overlapping with primers on average (3.67 vs. 18), as well as lower average frequency of those SNPs (3.03% vs. 10.56%) (Figure 4e). We also predicted the number of non-specific amplicons with BLAST, and Olivar had fewer non-specific amplicons on average compared to PrimalScheme (5 vs. 27, Figure 4f). In addition, the PrimalScheme design has 10 gaps (regions not covered by amplicon inserts, except for the left and right ends of the genome), with total length of 427nt (Supplementary Figure S1), while there were no gaps in any of the 6 Olivar designs. Default parameters of Olivar and PrimalScheme, SNP calling with Variant Database, generation of pseudo genomes for PrimalScheme and prediction of non-specific amplicons can be found in Methods.

**Figure 4.**
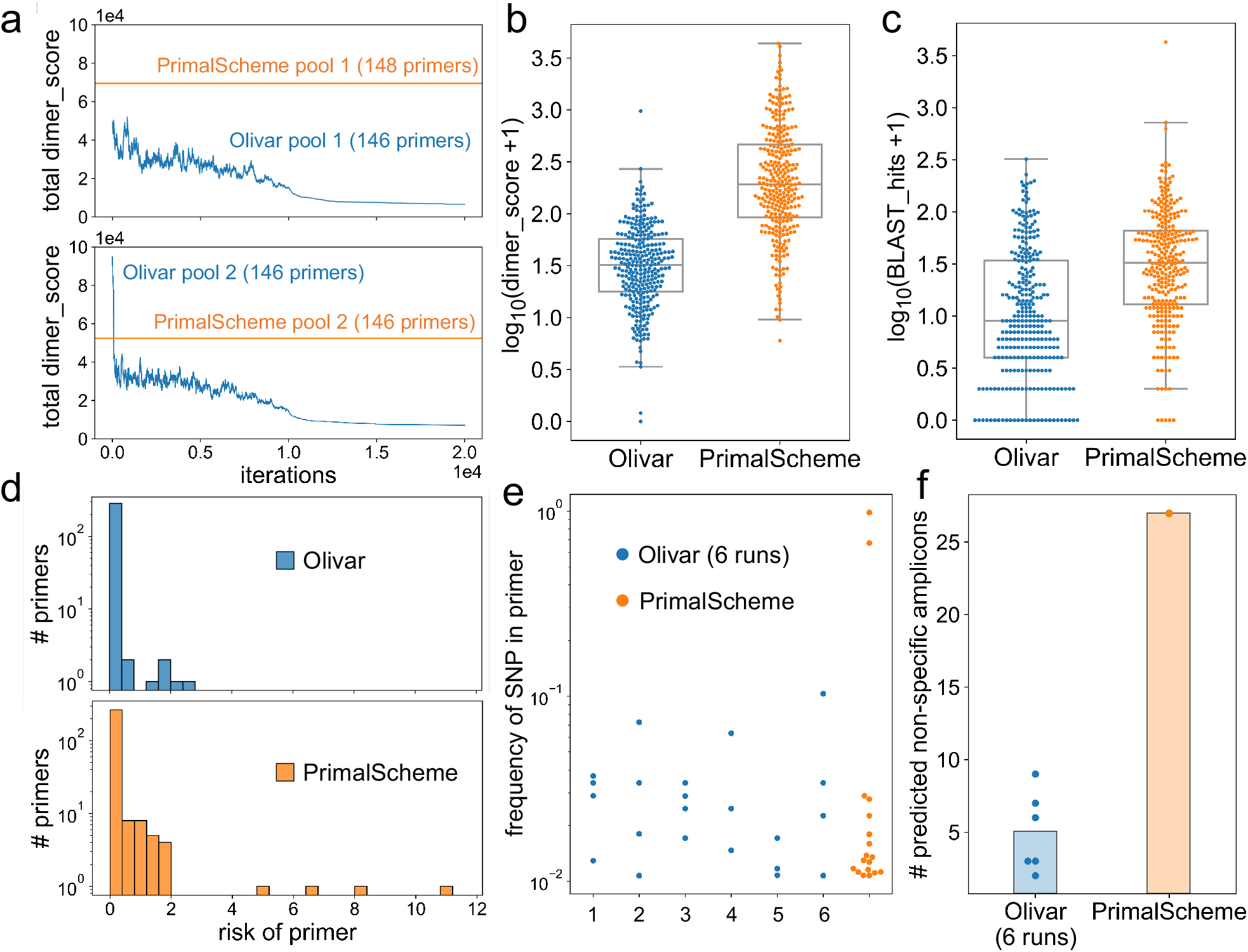
In silico comparison between Olivar and PrimalScheme on SARS-CoV-2 genome. **(a)** For each primer pool, primer dimerization is optimized with the previously published algorithm SADDLE. SADDLE calculates a dimer score for each primer within a primer pool, as an estimation of primer-primer interaction. Olivar pool 1 has an initial total dimer score of 49,692, and a final score of 6,640. Olivar pool 2 has an initial total dimer score of 95,747, and a final score of 7,192. PrimalScheme pool 1 and 2 have total dimer scores of 69,474 and 52,369, respectively. **(b)** The log10(dimer_score + 1) is calculated for each primer of Olivar or PrimalScheme (*p ≤*0.001, pool 1 and 2 are shown together). **(c)** The number of hits for each primer is acquired with the BLAST database of non-targeted sequences. Here only human genome is included. log10(BLAST_hits + 1) is calculated for each primer of Olivar or PrimalScheme (*p ≤*0.001, pool 1 and 2 are shown together). **(d)** The risk of a primer is the sum of all risk scores within the primer. Risk distribution is shown for all primers of Olivar or PrimalScheme. Olivar primers have average risk of 0.14, and PrimalScheme primers have average risk of 0.29. **(e**,**f)** Results of six Olivar runs with the same settings but different random seed. **(e)** Frequencies of the SNPs that overlap with primers. Out of 440 input SNPs, Olivar primers overlap with 3.67 SNPs on average, with average frequency of 3.03% and highest frequency of 10.72%, while PrimalScheme primers overlap with 18 SNPs with average frequency of 10.56% and highest frequency of 98.31%. **(f)** Six Olivar designs have 5 predicted non-specific amplicons on average, and the PrimalScheme design has 27 predicted non-specific amplicons. Details about the prediction of non-specific amplicons can be found in Methods.

Sequences and coordinates of both the Olivar design and the PrimalScheme design can be found in Supplementary Dataset.

### Experimental validation of Olivar on synthetic RNA

To further demonstrate Olivar’s performance in real-world applications, we ordered one of the Olivar-designed primer set described above (Figure 4a-d) and compared Illumina sequencing results with the widely used SARS-CoV-2 primer set ARTIC v4.1. Note that ARTIC v4.1 has additional primers added to target the Omicron variants and certain primers have double concentration in the primer pool for better coverage uniformity. We first tested both primer sets on synthetic SARS-CoV-2 RNA samples (Twist Bioscience), with different RNA concentrations, as determined by the Ct value from quantitative real-time PCR (qPCR) targeting the N gene. For both low Ct and high Ct samples, Olivar and ARTIC v4.1 had a similar percentage of sequencing reads mapped to the reference, ranging from 75% to 95%, while Olivar had a lower amount of bases with less than 0.05× median coverage (Table 1). Detailed description of experimental protocols and analysis of sequencing results can be found in Methods.

**Table 1.**
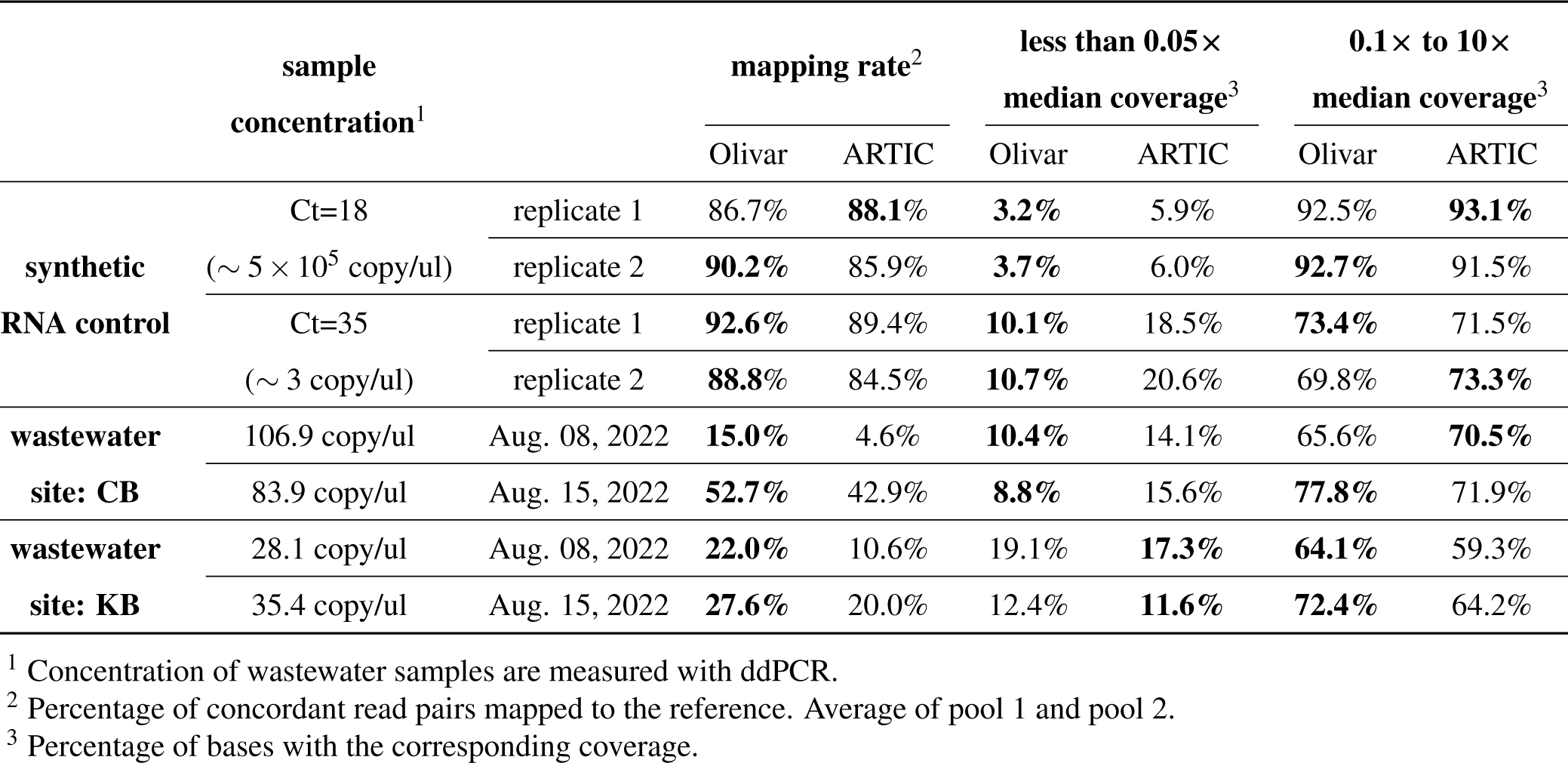
Mapping rate and coverage uniformity of Olivar Sars-Cov-2 primers and ARTIC v4.1 primers.

### Sequencing SARS-CoV-2 from wastewater with Olivar primers and ARTIC v4.1

Targeted amplification of pathogen genomes from wastewater samples is challenging since the targeted genomes are highly fragmented, dilute, and comprised of mixtures of circulating variants^6^. We collected 4 wastewater samples from two locations in Houston, USA at two time points. Using the same primer sets and experimental protocol as above, we observed 1 to 3-fold higher mapping rates of Olivar than ARTIC v4.1, shown in Table 1. Olivar also had lower or similar percentage of low coverage bases, compared with ARTIC v4.1 (Table 1). Figure 5a shows the overall genomic coverage of both Olivar and ARTIC v4.1 for one of the wastewater samples (site: CB, Aug. 15, 2022), with amplicon locations shown in gray lines. To compare the coverage uniformity of Olivar and ARTIC v4.1, genomic locations in Figure 5a are sorted by coverage (Figure 5b), showing Olivar has fewer bases with low coverage (8.8% vs. 15.6%). This is likely due to Olivar designs having shorter amplicon lengths and more overlapping of amplicons since there is a smaller difference for low coverage amplicons (13.7% and 16.2% for Olivar and ARTIC v4.1, respectively), shown in Figure 5c. Coverage of other samples are shown in Supplementary Figure S2∼S8. Details about coverage calculation can be found in Methods.

**Figure 5.**
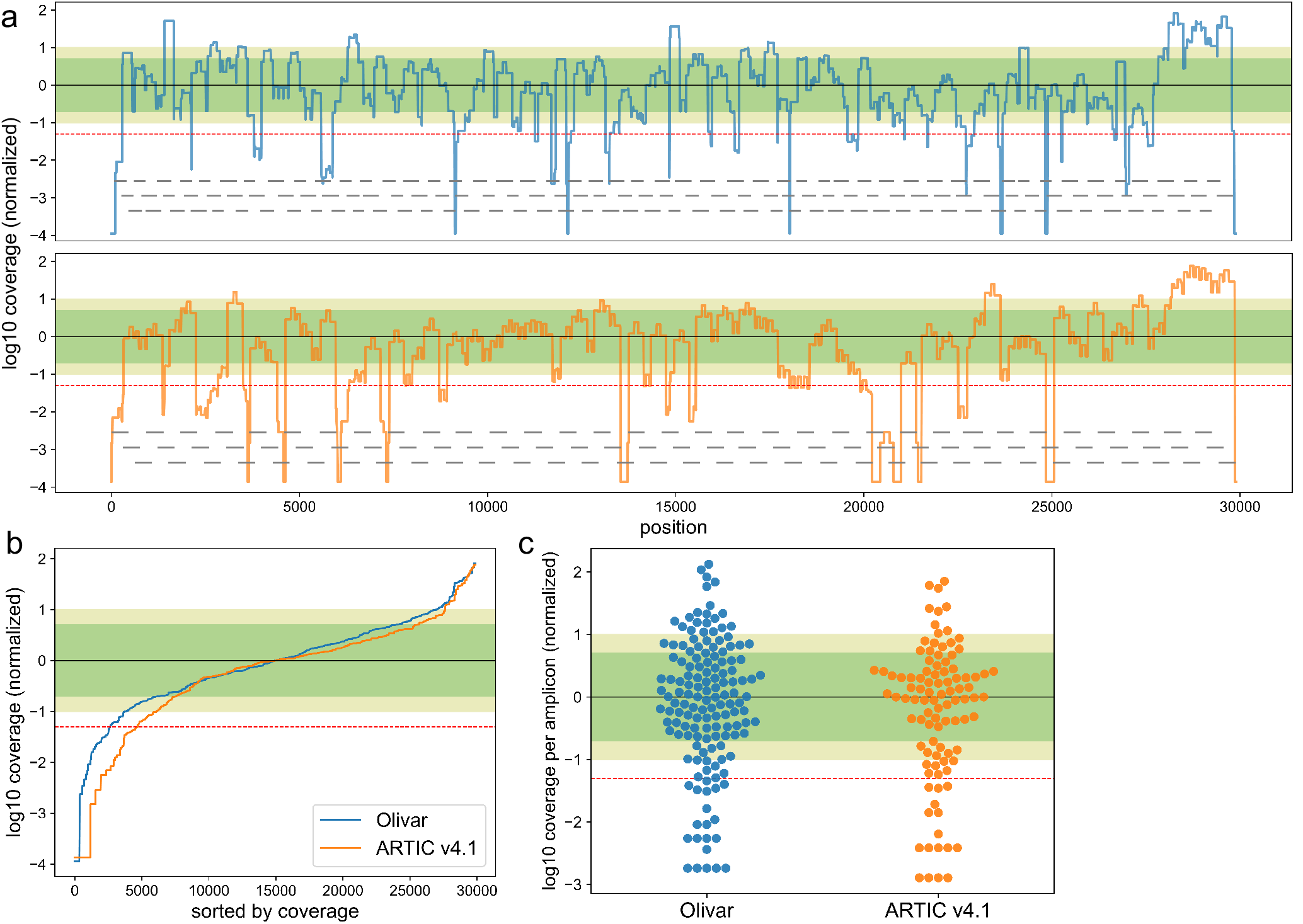
Sars-Cov-2 whole genome coverage of both Olivar (blue) and ARTIC v4.1 (orange) primers. Figures showing results from one wastewater sample (site: CB, Aug. 15, 2022). **(a)** log10 coverage of each base. Coverage is normalized by median coverage of all bases. Gray lines represent location of amplicons. **(b)** Sorted log10 coverage of each base. Black solid line represents the median coverage, green shade represents 0.2× to 5× median coverage (Olivar: 58.8% bases, ARTIC v4.1: 59.4% bases), olive shade represents 0.1 ×to 10× coverage (Olivar: 77.8% bases, ARTIC v4.1: 71.9% bases), red dashed line represents 0.05× median coverage (Olivar: 8.8% bases less than 0.05×, ARTIC v4.1: 15.6% bases less than 0.05×). Coverage uniformity of other samples is shown in Table 1. **(c)** log10 coverage of each amplicon, normalized by median amplicon coverage. (Olivar: 52.7% between 0.2× to 5× coverage, 67.8% between 0.1× to 10× coverage, 13.7% less than 0.05× coverage; ARTIC: 52.5% between 0.2× to 5× coverage, 67.7% between 0.1× to 10× coverage, 16.2% less than 0.05× coverage).

### Olivar web application

To enable easy access for researchers world wide, we developed a user friendly interactive web interface to run Olivar online (Figure 6). Although it does not support all available functions at the moment, design output is compatible with the local version of Olivar for further analysis. Input files and parameters are the same as the local version of Olivar, with preset defaults parameters and example input files for download. Each run is assigned with a unique identifier for downloading design results at any time.

**Figure 6.**
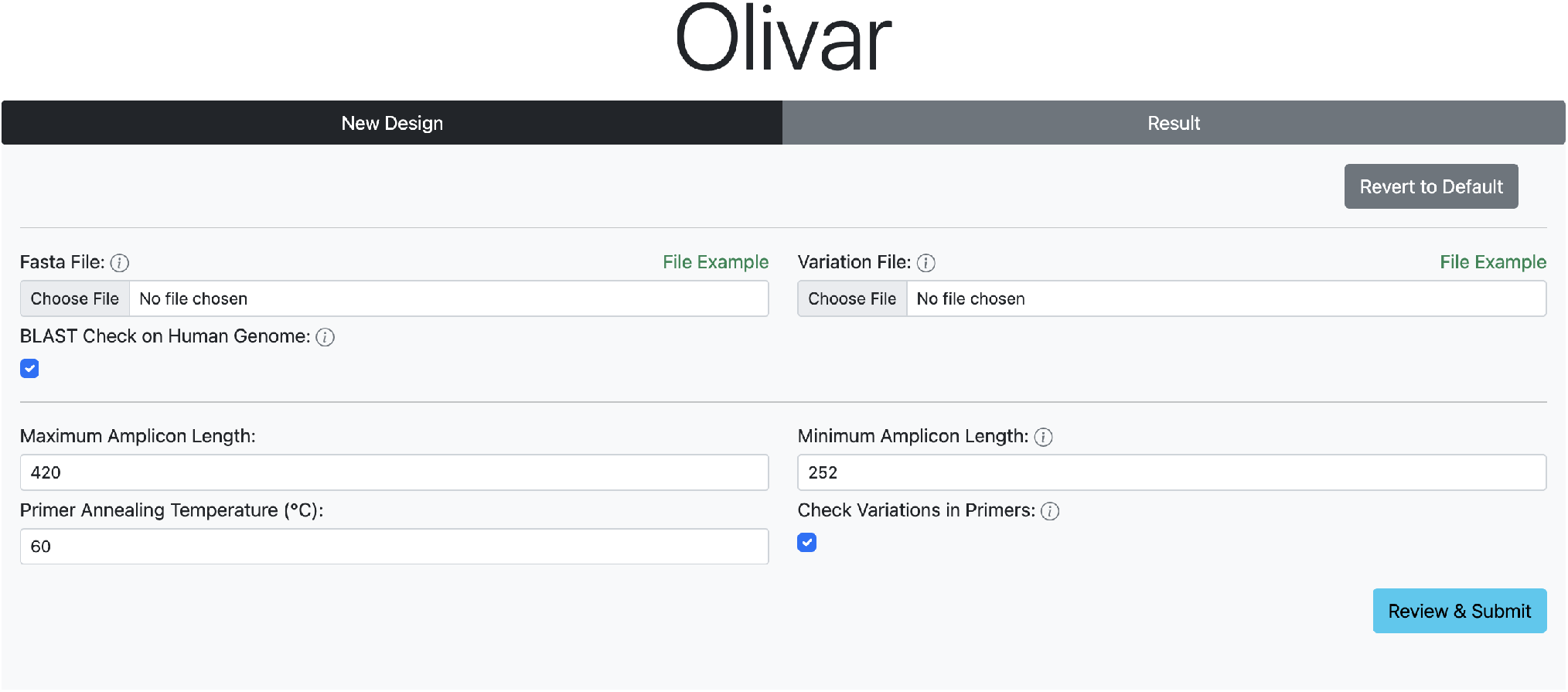
User interface of the Olivar web app. Users need to upload a reference sequence, and an optional list of SNP locations and frequencies. Other parameters can be adjusted according to application or kept as defaults.

## Discussion

We have described and presented results on Olivar, an approach designed towards a fully automated, end-to-end computational method for tiled primer design. Olivar quantifies undesired sequence features with single nucleotide risk scores across the whole genome, enabling efficient evaluation and optimization of primer design regions and ultimately improving the performance of output primers. In-silico validation shows that Olivar can avoid high-risk regions in the targeted genome, including SNPs, extreme GC content, low sequence complexity, and repetitive sequences. Compared with the state-of-the-art PCR tiling design software PrimalScheme, Olivar has fewer SNPs overlapping with primer and less predicted primer dimer and non-specific priming. In addition, in a head-to-head comparison with the most commonly used SARS-CoV-2 primer set ARTIC v4.1, Olivar offers equivalent to higher mapping rates and similar genomic coverage on both synthetic RNA samples and wastewater samples. These mapping rates and coverage improvements highlight that Olivar can provide robust designs capable of being produced at lower sequencing costs. Moreover, the ARTIC primer set has undergone several versions of optimization whereas the experimentally validated Olivar primers were automatically designed, saving significant time and cost that comes with multiple rounds of manual redesign.

Furthermore, Olivar is a versatile approach not limited to designing tiled amplicons for viral genomes. For example, PCR tiling is frequently used in applications such as full-length sequencing of entire genes^21^. An Olivar risk array could not only guide the design of tiled amplicons, but also one or a few amplicons when there is no strict requirement on amplicon location (e.g., pathogen detection with digital PCR^6^ and measuring copy number variation^22^), by selecting genomic regions with low-risk scores. This could help users quickly find signature sequences ideal for downstream design, such as non-repetitive and highly conserved regions. Furthermore, the modular nature of calculating a risk array allows the customization of risk components. For example, in applications where SNPs are needed to distinguish similar species or strains of pathogens^23^, users could define risk scores for sensitivity and specificity to find sequences targeting desired strains while avoiding unwanted ones.

Despite the automatic design workflow of Olivar, user’s input of sequence variation information and a BLAST database of background non-specific sequences are needed to describe the design problem sufficiently. For sequence variation, it is represented with a list of locations and frequencies and input to Olivar. Another way to represent sequence variation is directly using the group of genomic sequences to be covered (e.g, strains of a certain virus)^4^. While the latter is usually more readily available than the former for viruses and bacteria, it needs more computational resources to evaluate the sensitivity of a PDR for a group of genomes through local alignment or multiple sequence alignment (MSA), especially when the number of genomes is large. For well-studied pathogens, such as SARS-CoV-2 and human monkeypox virus, the MSA Olivar uses is updated in real-time from the public database GISAID^9^ and coordinates and frequencies of nucleotide change are available at Nextstrain^19^. For species without publicly available MSA, tools such as Parsnp^24^ can efficiently build core-genome alignment and make variant calls.

Another user-defined input for Olivar is the BLAST database for background non-targeted sequences. While we provide the BLAST database for the human genome as it is frequently considered background, users will likely need more comprehensive and application-specific databases to reduce non-specific byproducts further. Leveraging GC content and sequence complexity can also help avoid low-complexity sequences. We are actively building more background databases for various scenarios, such as pathogen detection in wastewater. Additional background sequence screening improvements will be included in future updates of Olivar.

Except for user input data, parameters in the design workflow could also affect the output primer set. The parameters in Olivar workflow can be categorized into two groups: 1) parameters in risk array calculation, and 2) parameters in the optimization phase, including PDR optimization and primer dimer optimization. The first group of parameters are set based on the fact that most PCR tiling assays are run under annealing temperature of 50°C-70°C, with 55°C-65°C being the most frequent setting. That results in primer length of 18nt-35nt, and we determined PDR length of 40nt to ensure there are enough primer candidates, while limiting the uncertainty of the final position of the primer. The generation of the risk array and its parameters are then determined based on the length of PDR (e.g, word size of 28, sequence complexity threshold of 0.4, and BLAST parameters, etc., with details described in the Method section). We also introduce weights for different risk components to allow more flexibility for users with expertise and have specific requirements for their design. Within the second group of parameters, parameters related to primer dimer optimization are well established and are adjusted automatically based on the size of the genome^11^. For PDR optimization, we investigated the level of greediness introduced in selecting PDR candidates and chose the optimal parameter based on in-silico experiments, as shown in Figure 2.

To our knowledge, Olivar is the first open-source computational tool designed to allow fully automatic design of multiplexed PCR tiling assays while considering SNPs, primer dimers, and non-specific amplification simultaneously, with the potential of significantly reducing manual redesign while maintaining low cost. We anticipate that Olivar will aid the surveillance of future infectious disease outbreaks by providing an automated tool for designing tiled amplicons.

## Methods

### Generation of risk array

A risk array is an array of non-negative real numbers

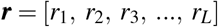

where *L* is the length of the targeted sequence. The array ***r*** consists of four weighted components,

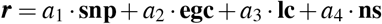

where *a*_1_, *a*_2_, *a*_3_ and *a*_4_ are set to 1 as default. **snp** represent SNPs, **egc** represent extreme GC content, **lc** represent low sequence complexity and **ns** represent non-specificity. For each user provided single nucleotide polymorphism (SNP) at position p of the targeted sequence,

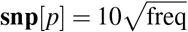

where freq could be the frequency of the SNP to be avoided. The root of SNP frequency is taken to amplify low frequency SNPs, and timed by a factor of 10 to be comparable to other risk components. Note that freq could also be user defined (e.g., when certain SNPs are considered more important than others, or simply just scale up all frequencies by a certain factor). To calculate the other three components, a set of equal-length, overlapping, and evenly distributed “words” are generated from the targeted sequence. Suppose the length of each word is ws (28 by default), *L* is divisible by ws and ws is divisible by a positive integer *c* (2 by default), then the distance between neighboring words is ws*/c*, and

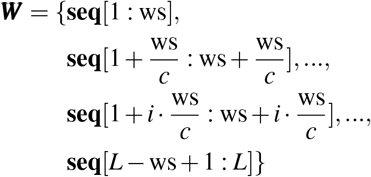

where ***W*** is the set of words, **seq** is the targeted sequence and *i* is a positive integer. If *L* is not divisible by ws, the targeted sequence is minimally trimmed to satisfy this condition, and corresponding elements in risk array ***r*** are set to 0. The value of ws is set as 28 by default based on typical primer length (20-25bp) and the minimum word size (7) of the NCBI BLAST+ program^25^. *c* is set as 2 by default so that each base is covered by two words. GC content and sequence complexity of each word is calculated. Sequence complexity is calculated based on Shannon entropy (described below). The number of BLAST hits of each word is acquired with the user-provided BLAST database. The following parameters are set to the NCBI BLAST+ program^25^ (version 2.12.0): e-value as 5, reward as 1, penalty as −3, gapopen as 5, gapextend as 2. These are the default parameters of the task “blastn-short”. For a nucleotide at position 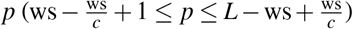 of the targeted sequence (except for head and tail regions), there is a set of *c* words overlapping with that nucleotide, denoted as ***W*** _***p***_ ⊂***W***. The average GC content of ***W*** _***p***_, the average sequence complexity of ***W*** _***p***_ and the average number of BLAST hits of ***W*** _***p***_ are denoted as gc, cmplx and hits, respectively. **nsraw** is an array of 0s with length of *L*. Then,

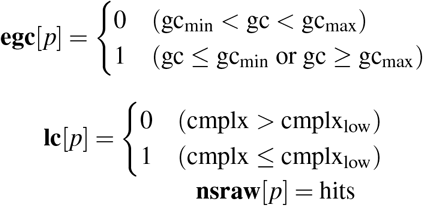

where gc_min_ (0.25 by default), gc_max_ (0.75 by default) and cmplx_low_ (0.4 by default) are user defined. **nsraw** is then normalized to get **ns**

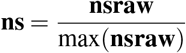

#### Calculation of sequence complexity

Sequence complexity is calculated based on Shannon’s entropy. For a DNA sequence ***S*** of length *L* (*L >* 3) with alphabet {A, T, C, G}, the number of each *k*-mer (*k* = 1, 2, 3) is stored as an array

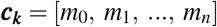

where *m* is the number of a certain k-mer, and *n ≤*4^*k*^. For example, if ***S*** = ACGCAGCGAGCAG, then ***c***_**1**_ = [4, 4, 5] since there are 4 ‘A’s, 4 ‘C’s and 5 ‘G’s. Then,

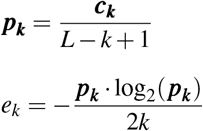

where *e*_*k*_ is the dot product of the two vectors. The complexity of ***S*** is the smallest of *e*_1_, *e*_2_ and *e*_3_.

### Optimization of PDRs

A PDR is defined by its start coordinate *p* and stop coordinate *p* + *l* − 1 (closed interval), where *l* is the length of the PDR (40 by default). All PDRs have the same length. The risk of the PDR is then defined as sum(***r***[*p* : *p* + *l −*1]), where ***r*** is the risk array.

#### Generation of one PDR

Each PDR is selected within a certain region ***C***, the length of which is greater than or equal to *l*. The risk of all possible PDRs within ***C*** is calculated, and PDRs with risk below *X* th percentile are considered candidate PDRs. Here percentile is defined as a PDR risk threshold below which percentage *X* of PDRs fall. Therefore, a lower *X* means a more stringent selection on PDR candidates. *X* is set to 30 by default.

#### Generation of a set of PDRs

PDRs are regions where primer candidates are generated, and primer candidates are subsequences of the PDR. Similar to forward primer (fP) and reverse primer (rP), there are also forward PDR (fPDR) and reverse PDR (rPDR). There are three restrictions for a valid PDR set given a targeted sequence:

1. PDRs should not overlap with each other (this guarantees that primers will not overlap).
2. A PDR pair should be able to generate amplicons that satisfy desired amplicon length, as defined by the user.
3. The targeted sequence is fully covered with inserts. An insert is the region between a PDR pair.

Suppose a valid PDR set consists of *N* PDR pairs and each PDR pair is generated sequentially. For the *k*th PDR pair (*k* ≥ 1), the fPDR is selected from region ***C***_2*k*−1_, and the coordinates of the fPDR is [*p*_2*k*−1_, *p*_2*k*−1_ + *l −*1]. The rPDR is selected from region ***C***_2*k*_, and the coordinates of the rPDR is [*p*_2*k*_, *p*_2*k*_ + *l −*1]. ALEN_min_ is minimum amplicon length and ALEN_max_ is maximum amplicon length.

Generation of a PDR set starts with the first PDR pair (*k* = 1). Starting from the left end of the risk array and the targeted sequence, the fPDR of the first PDR pair is selected from region ***C***_**1**_,

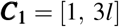

Here ***C***_**1**_ starts at 1 for simplicity. There are 2*l* + 1 possible PDRs within ***C***_**1**_, and one of them is selected as the first fPDR, with start coordinate *p*_1_, based on the method described in the section above (Generation of one PDR). The first rPDR is selected from region ***C***_**2**_, and based on the amplicon length restriction,

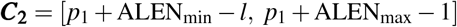

The start coordinate of the rPDR is selected as *p*_2_.

For the second PDR pair (*k* = 2), the fPDR and rPDR are selected from region ***C***_**3**_ and ***C***_**4**_, respectively. Considering the three restrictions, we have

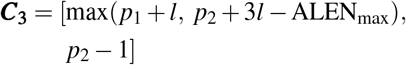

The second fPDR is selected from ***C***_**3**_ with start coordinate *p*_3_. Then,

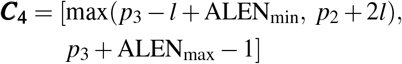

The second rPDR is selected from ***C***_**4**_ with start coordinate *p*_4_.

Starting from the third PDR pair (*k≥* 3), the fPDR and rPDR are selected from region ***C***_2*k*−1_ and ***C***_2*k*_, respectively. Considering the three restrictions, we have

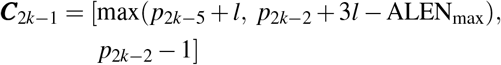

The *k*th fPDR is selected from ***C***_2*k*−1_ with start coordinate *p*_2*k*−1_. Then,

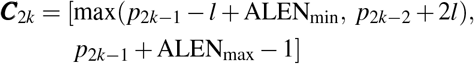

The *k*th rPDR is selected from ***C***_2*k*_ with start coordinate *p*_2*k*_.

PDR generation will stop when ***C***_2*k*_ exceeds the limit of the targeted sequence.

#### Loss of a PDR set

The Loss of a PDR set is defined as the total risk of the top 10% high-risk PDRs. For an input genome of length *L*, the number of PDR sets generated is ⌊500*L/*ALEN_max_⌋. The PDR set with the lowest Loss is selected for downstream design.

#### Primer pool assignment

If the targeted sequence is fully covered with inserts (region between fP and rP), then amplicons must be overlapping and the *k*th fP could form a short amplicon with the (*k −*1)th rP. These short amplicons are usually much shorter than desired amplicons, having much higher amplification efficiency. To avoid the formation of these short amplicons, most PCR tiling applications assign amplicons into two pools so that amplicons within each pool are non-overlapping. Suppose the first PDR pair is assigned to pool 1 and the second PDR pair is assigned to pool 2, then the 2*k* − 1the PDR pair is assigned to pool 1 and the 2*k*th PDR pair is assigned to pool 2 (*k* ≥ 2).

### Optimizing primer candidates with SADDLE

In the previous step the optimal PDR set is separated into two pools of PDRs, and each pool is input to SADDLE separately. Primer candidates are first generated for each PDR, with default parameters: temperature as 60°C, salinity as 0.18, max primer length as 36, max Δ*G* (Gibbs free energy) as −11.8. One primer candidate is randomly picked from each PDR to form a primer set. SADDLE minimize primer dimers within the randomly generated primer set by iteratively replacing a primer candidate with another one from the same PDR. The number of iterations is determined by the total number of PDRs in the pool. The final optimized primer set will be the output of Olivar.

### Prediction of non-specific amplicons

The prediction of non-specific amplicons starts with running BLAST through each single primer, with the following parameters set to the NCBI BLAST+ program^25^: evalue as 5000, reward as 1, penalty as −1, gapopen as 2, gapextend as 1. The values of these parameters are determined based on Primer BLAST^26^. For each BLAST hit, the BLAST program outputs the location and orientation of that hit. A single primer could have multiple BLAST hits, distributed to different chromosomes. For each chromosome, the location and orientation of hits of all primers are analyzed together. If two hits are close enough and the orientations are legitimate for PCR, that pair of hits is reported as a predicted non-specific amplicon.

### Calculation of Nextstrain nucleotide entropy

Nextstrain^19^ calculates Shannon’s entropy for each nucleotide position of the reference SARS-CoV-2 genome based on MSA of genomes available on GISAID^9^. Here we include global genomes date from Dec. 21, 2019 to Dec. 04, 2022. Entropy at a given position *c* is calculated as below. Suppose we have *N* genomes. At position *c, N*_1_ genomes has ‘A’, *N*_2_ genomes has ‘T’, *N*_3_ genomes has ‘C’, *N*_4_ genomes has ‘G’. Suppose *N*_1_, *N*_2_, *N*_3_ and *N*_4_ are non-zero.

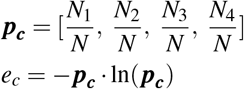

where *p*_*c*_ is an array and *e*_*c*_ is dot product and entropy at position *c*. Of 7,774 locations with non-zero entropy, we selected 1,517 locations with entropy greater than 0.01 and input to Olivar as SNP locations and frequencies. We used the reference genome provided by Nextstrain (GenBank MN908947.3) without a BLAST database of non-targeted sequences. Other parameters are kept as default.

### SNP calling with Variant Database

We downloaded MSA and metadata of GISAID SARS-Cov-2 genomes before Feb. 28, 2022 and used Variant Database^20^ (version 2.4) to output SNPs of lineage B.1.617.2 and B.1.1.529, with GISAID genome EPI_ISL_402124 as reference for coordinates. Variant Database compares the differences between reference genome EPI_ISL_402124 and all genomes in lineage B.1.617.2 (or B.1.1.529), and SNP frequency is defined as the percentage of genomes that are different from the reference at a given location. Note that Variant Database version 2.4 does not output insertions, so here SNPs include substitutions and deletions. We used the command “frequencies <cluster>” to list the frequencies of individual mutations for B.1.617.2 and B.1.1.529 separately. Insertions are not generated due to the limitations of the Variant Database. Of the 10,000 output SNPs, we kept 440 SNPs with a frequency greater than 0.01 and input to Olivar.

### In silico comparison of Olivar and PrimalScheme

For Olivar input, we used GISAID genome EPI_ISL_402124 as a reference, a BLAST database of human genome assembly GRCh38.p13 as non-targeted sequences, and 440 SNPs of SARS-CoV-2 lineage B.1.617.2 and B.1.1.529 generated with Variant Database (described above). For Olivar and PrimalScheme (version 1.4.1), desired amplicon length is set to 252nt to 420 nt. Other Olivar and PrimalScheme parameters are kept as default.

#### Input genomes for PrimalScheme

100 artificial genomes are created, bearing the same 440 SNPs input to Olivar. Each SNP is randomly put back to *n* of the 100 copies of the GISAID reference EPI_ISL_402124,

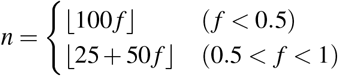

where *f* is the frequency of the SNP. We did not keep the original frequency for high-frequency SNPs in the artificial genomes because PrimalScheme will consider a position to be conserved if the vast majority of genomes have the same SNP and that position will not be avoided.

### Sample preparation and quantification

Synthetic RNA control for SARS-CoV-2 is purchased from Twist Bioscience (part number 105204, GISAID ID EPI_ISL_6841980). Time-weighted composite samples of raw wastewater were collected every 1h for 24 h from the influent of the two domestic wastewater treatment plants (WWTPs), CB and KB, on two separate dates (Aug. 8 and Aug. 15, 2022). Samples were kept on ice during transport and stored at 4°C in the laboratory to be processed within 24 hours of collection. Sample concentration using HA filtration and bead beating methods, as well as RNA extraction method using the ChemagicTM Prime Viral DNA/RNA 300 Kit H96 (Chemagic, CMG-1433, PerkinElmer) were as decribed in ref.^6^. To normalize synthetic RNA control to Ct values of 18 and 35, One-step RT-qPCR were performed with qPCRBIO probe 1-Step Go LoROX (PB25.41, PCR Biosystems) on the QuantStudio 3 Real Time PCR System (A28567, Applied Biosystems) as previously described in ref.^27^. These two Ct bounds were selected to represent the high and low concentrations of SARS-CoV-2. SARS-CoV-2 concentrations of wastewater samples were quantified using RT-ddPCR on a QX200 AutoDG Droplet Digital PCR System (Bio-Rad) and a C1000 Thermal Cycler (Bio-Rad) in 96-well optical plates, as previously described in ref.^6^.

### Multiplexed PCR, library preparation and sequencing

ARTIC V4.1 primer panel was purchased from IDT (Artic V4.1 NCOV-2019 Panel, 500rxn, 10011442). Olivar primers were ordered in tubes from Sigma Aldrich and mixed by hand to achieve the final concentration of 15 nanomolar (nM) per primer. Reverse transcription of synthetic RNA control and extracted wastewater RNA were conducted using 8 uL of sample RNA and LunaScript RT SuperMix kit (NEB, E3010), as described in ref.^17^. To avoid bias attributed to reverse transcription, for each sample, the total volume of 10 uL cDNA product were gently homogenized by pipetting then divided into four 2.5 uL aliquots for the downstream PCR amplification reactions (using primer pool 1 and 2 of ARTIC V4.1, and using primer pool 1 and 2 of Olivar). PCR amplification was also performed using Q5 Hot Start High-Fidelity 2X Master Mix (NEB M0494) as described in ref.^17^. PCR products were purified using AmPureTM XP beads (Beckman Coulter Inc., A63880). A high bead-to-sample ratio of 1.8 was applied to maximize the potentials of capturing PCR byproduct. Purified DNA samples were normalized to 20 ng/uL in 25 uL and submitted for amplicon sequencing service at Azenta (EZ-Amplicon), with paired-end (2 ×250bp), adapter trimmed Illumina reads output. The concentration of amplicons was measured using Qubit dsDNA HS kit and a Qubit 2.0 fluorometer (Invitrogen).

### Analysis of sequencing data

Paired-end sequencing reads are mapped to the reference genome (GISAID ID EPI_ISL_402124) with Bowtie2 (version 2.4.5)^28^, with maximum fragment length for valid paired-end alignments (-X) set to 1000. Coverage of each reference position is calculated as the number of reads overlapping that position, using PySAM (version 0.19.1)^29^. Sequencing reads are also mapped to amplicon sequences with Bowtie2, and the coverage for each amplicon is the number of read pairs mapped to that amplicon.

## Acknowledgements

The authors thank Nina G. Xie for experimental advice.

## Author contributions statement

M.X.W, L.B.S and T.J.T conceived the project. L.B.S and T.J.T supervised the project. M.X.W and E.G.L conducted the experiments. E.K developed the web app. M.X.W and B.K developed the software. All authors conceived the experiments and drafted the original paper. All authors read, revised, and approved the manuscript.

## Code Availability

Source code, installation guide and usage of Olivar are available at https://github.com/treangenlab/Olivar.

## Data Availability

Sequencing data in this study is available at NCBI SRA under BioProject PRJNA911448. Sequences and coordinates of primers, Nextstrain entropy, location and frequencies of SNPs, and mapping rates of sequencing samples are available in Supplementary Dataset (also available at https://github.com/treangenlab/Olivar).

