## Supplementary Information for "Olivar: automated variant aware primer design for multiplex tiled amplicon sequencing of pathogens"

### ABSTRACT

Tiled amplicon sequencing has served as an essential tool for tracking the spread and evolution of SARS-CoV-2 in real-time directly from environmental and clinical samples. Over 14 million SARS-CoV-2 genomes are now available on GISAID, most sequenced and assembled via tiled amplicon sequencing. While computational tools for tiled amplicon design exist, they require downstream manual optimization both computationally and experimentally, which is slow, laborious, and costly. Here, we present Olivar, the first open-source computational tool capable of fully automating the design of tiled amplicons by integrating SNPs, non-specific amplification, etc. into a "risk score" for each single nucleotide of the target genome. Olivar evaluates thousands sets of possible tiled amplicons and minimizes primer dimer in parallel. In a direct in-silico comparison with PrimalScheme, we show that Olivar has fewer SNPs overlapping with primers and predicted PCR byproducts. We also compared Olivar head-to-head with ARTIC v4.1, the most widely used tiled amplicons for SARS-CoV-2 sequencing. We next tested Olivar on real wastewater samples and found that our automated approach had up to 3-fold higher mapping rates compared to ARTIC v4.1 while retaining similar coverage. To the best of our knowledge, Olivar represents the first open-source, fully automated design tool that simultaneously evaluates and optimizes risks of known primer design issues for robust tiled amplicon sequencing. Olivar is available as a web application at `https://olivar.rice.edu/`. Olivar can also be installed locally as a command line tool with Bioconda. Source code, installation guide and usage are available at `https://gitlab.com/treangenlab/olivar`.

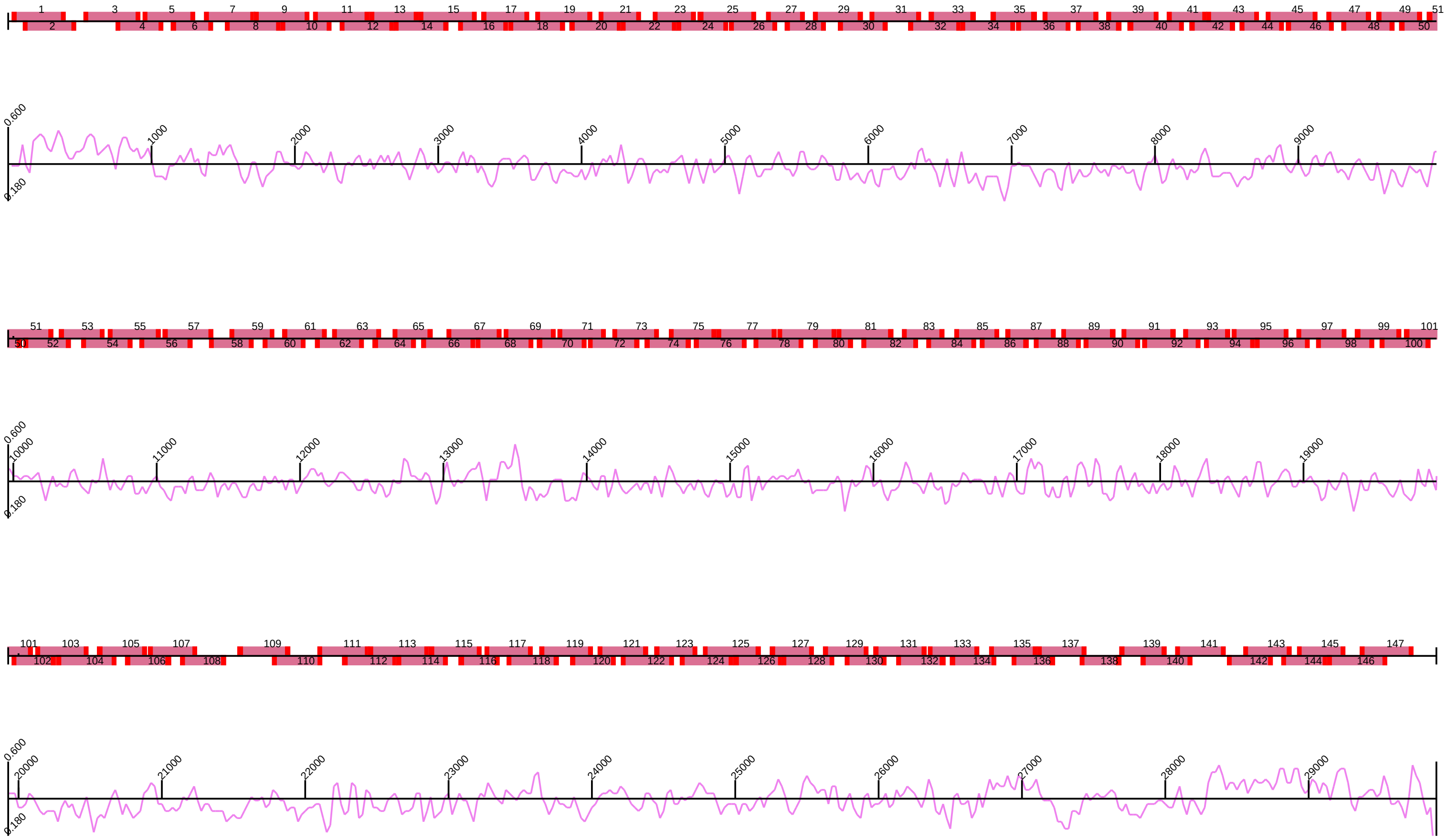


**Figure 1.** Location of amplicons designed by PrimalScheme, generated with the PrimalSheme software. 10 gaps are found in the PrimalScheme design: 448 to 553 (106bp), 9221 to 9225 (5bp), 11368 to 11393 (26bp), 12539 to 12539 (1bp), 21420 to 21561 (142bp), 22093 to 22113 (21bp), 26795 to 26799 (5bp), 27423 to 27431 (9bp), 27663 to 27710 (48bp) and 28395 to 28458 (64bp).

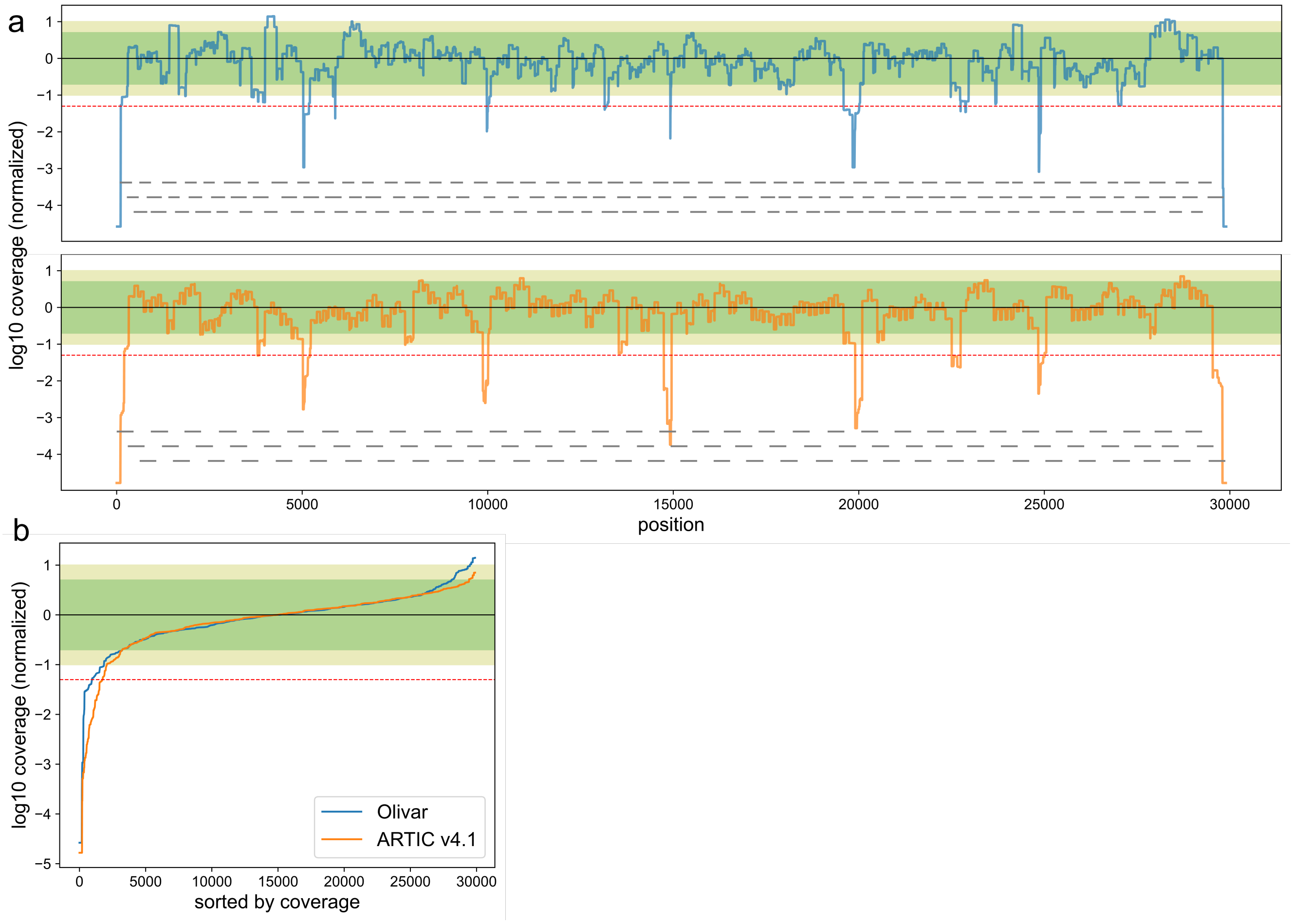


**Figure 2.** **Sars-Cov-2 whole genome coverage of both Olivar (blue) and ARTIC v4.1 (orange) primers. Figures showing results from one wastewater sample (Ct=18, replicate 1). (a)** log10 coverage of each base. Coverage is normalized by median coverage of all bases. Gray lines represent location of amplicons. **(b)** Sorted log10 coverage of each base. Black solid line represents the median coverage, green shade represents 0.2× to 5× median coverage (Olivar: 83.2% bases, ARTIC v4.1: 87.6% bases), olive shade represents 0.1× to 10× coverage (Olivar: 92.5% bases, ARTIC v4.1: 93.1% bases), red dashed line represents 0.05× median coverage (Olivar: 3.2% bases less than 0.05×, ARTIC v4.1: 5.9% bases less than 0.05×).

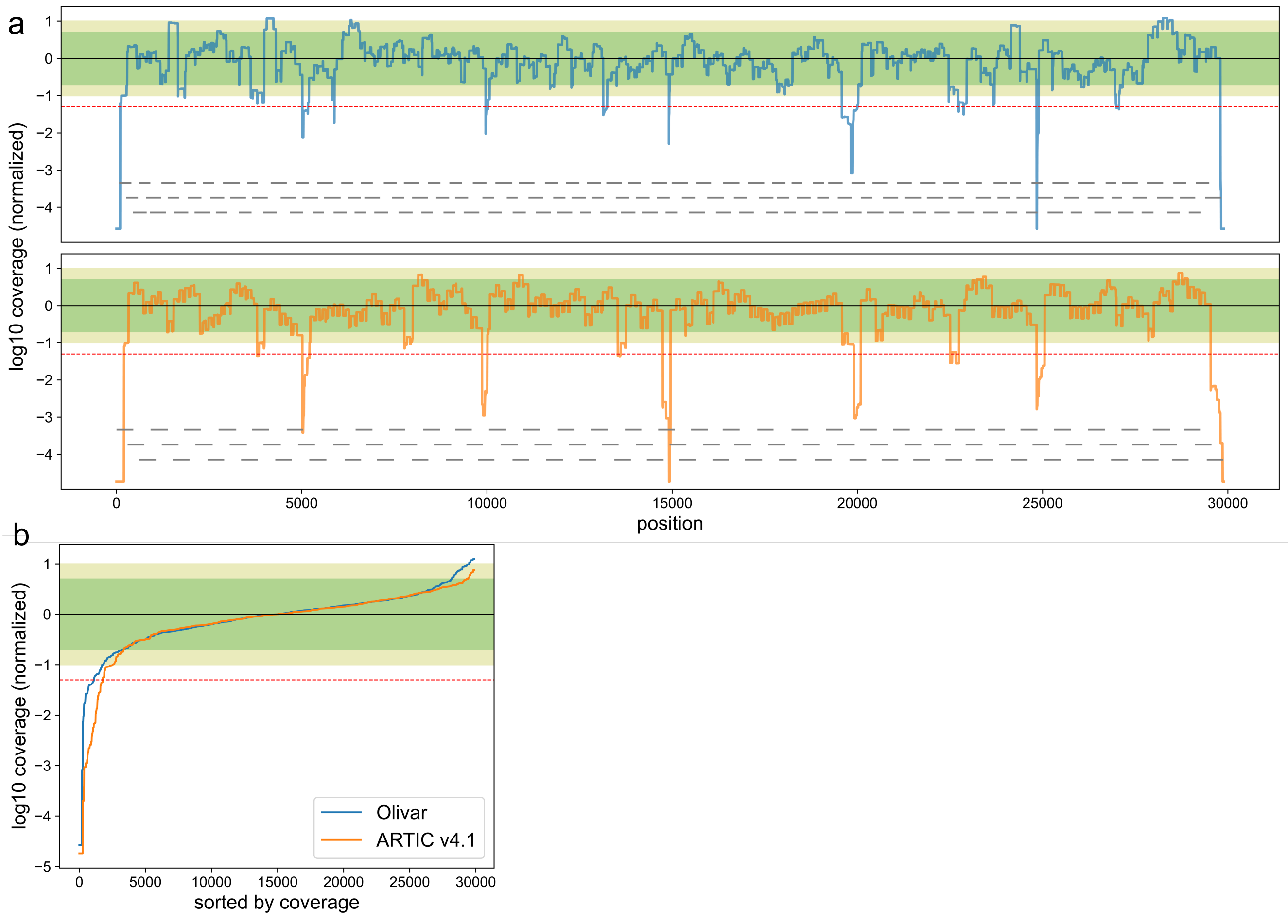


**Figure 3. Sars-Cov-2 whole genome coverage of both Olivar (blue) and ARTIC v4.1 (orange) primers. Figures showing results from one wastewater sample (Ct=18, replicate 2).** (**a**) log10 coverage of each base. Coverage is normalized by median coverage of all bases. Gray lines represent location of amplicons. **(b)** Sorted log10 coverage of each base. Black solid line represents the median coverage, green shade represents 0.2× to 5× median coverage (Olivar: 83.3% bases, ARTIC v4.1: 86.8% bases), olive shade represents 0.1× to 10× coverage (Olivar: 92.7% bases, ARTIC v4.1: 91.5% bases), red dashed line represents 0.05× median coverage (Olivar: 3.7% bases less than 0.05×, ARTIC v4.1: 6.0% bases less than 0.05×).

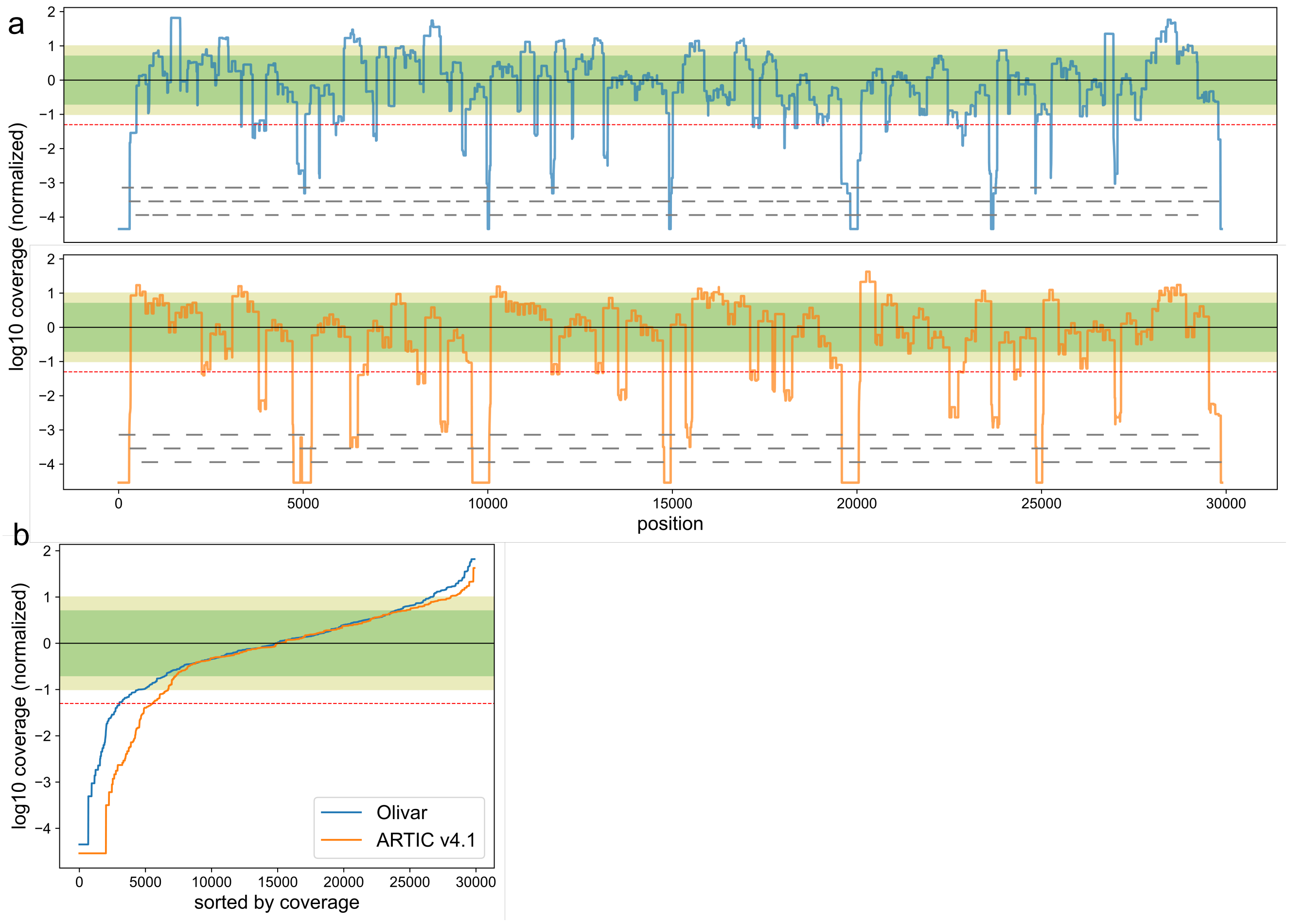


**Figure 4.** **Sars-Cov-2 whole genome coverage of both Olivar (blue) and ARTIC v4.1 (orange) primers. Figures showing results from one wastewater sample (Ct=35, replicate 1).** **(a)** log10 coverage of each base. Coverage is normalized by median coverage of all bases. Gray lines represent location of amplicons. **(b)** Sorted log10 coverage of each base. Black solid line represents the median coverage, green shade represents 0.2× to 5× median coverage (Olivar: 57.9% bases, ARTIC v4.1: 56.8% bases), olive shade represents 0.1× to 10× coverage (Olivar: 73.4% bases, ARTIC v4.1: 71.5% bases), red dashed line represents 0.05× median coverage (Olivar: 10.1% bases less than 0.05×, ARTIC v4.1: 18.5% bases less than 0.05×).

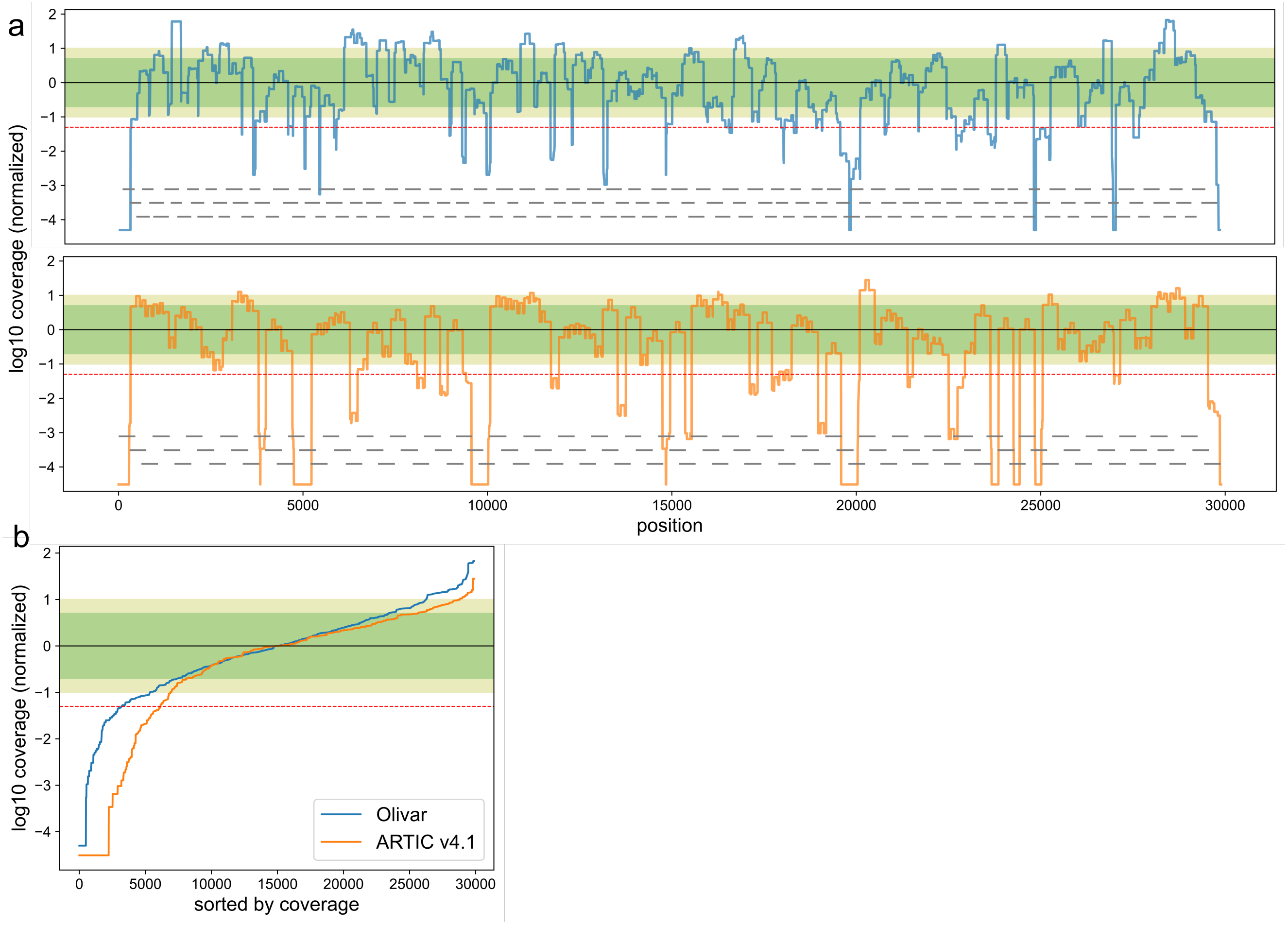


**Figure 5.** **Sars-Cov-2 whole genome coverage of both Olivar (blue) and ARTIC v4.1 (orange) primers. Figures showing results from one wastewater sample (Ct=35, replicate 2).** (**a**) log10 coverage of each base. Coverage is normalized by median coverage of all bases. Gray lines represent location of amplicons. **(b)** Sorted log10 coverage of each base. Black solid line represents the median coverage, green shade represents 0.2× to 5× median coverage (Olivar: 53.3% bases, ARTIC v4.1: 58.4% bases), olive shade represents 0.1× to 10× coverage (Olivar: 69.8% bases, ARTIC v4.1: 73.3% bases), red dashed line represents 0.05× median coverage (Olivar: 10.7% bases less than 0.05×, ARTIC v4.1: 20.6% bases less than 0.05×).

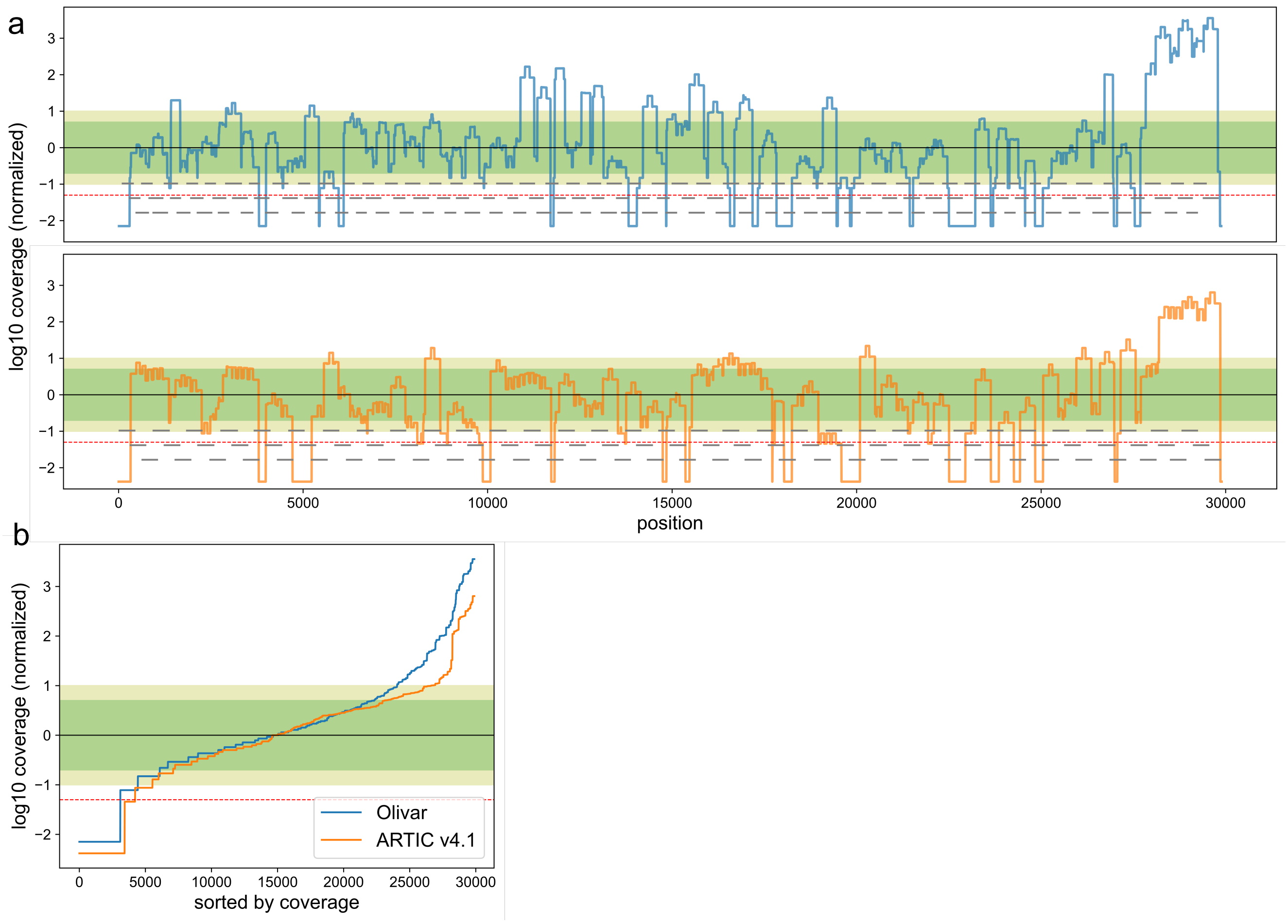


**Figure 6.** **Sars-Cov-2 whole genome coverage of both Olivar (blue) and ARTIC v4.1 (orange) primers. Figures showing results from one wastewater sample (site: CB, Aug. 08, 2022). (a)** log10 coverage of each base. Coverage is normalized by median coverage of all bases. Gray lines represent location of amplicons. **(b)** Sorted log10 coverage of each base. Black solid line represents the median coverage, green shade represents 0.2× to 5× median coverage (Olivar: 53.7% bases, ARTIC v4.1: 53.3% bases), olive shade represents 0.1× to 10× coverage (Olivar: 65.6% bases, ARTIC v4.1: 70.5% bases), red dashed line represents 0.05× median coverage (Olivar: 10.4% bases less than 0.05×, ARTIC v4.1: 14.1% bases less than 0.05×).

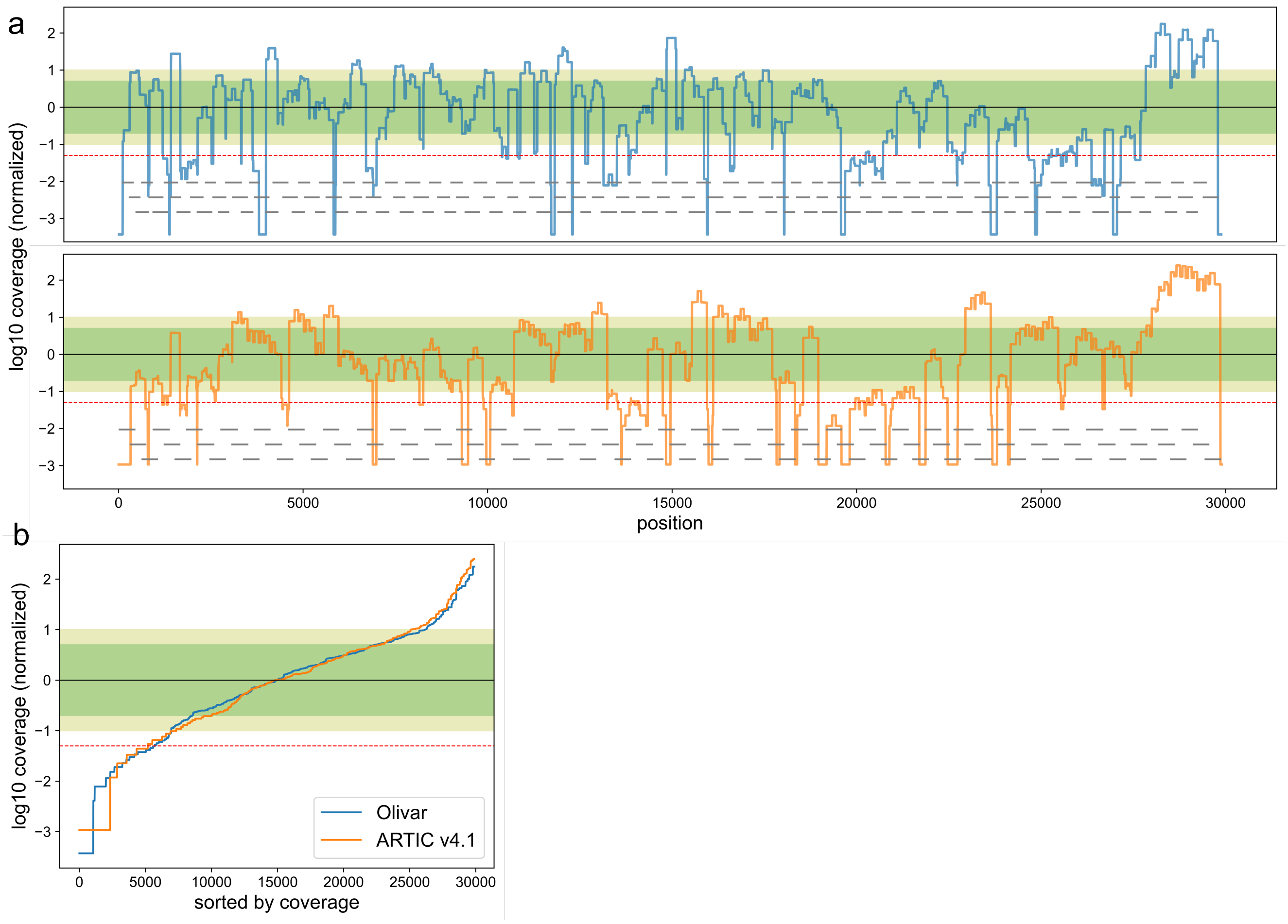


**Figure 7.** **Sars-Cov-2 whole genome coverage of both Olivar (blue) and ARTIC v4.1 (orange) primers. Figures showing results from one wastewater sample (site: KB, Aug. 08, 2022).** **(a)** log10 coverage of each base. Coverage is normalized by median coverage of all bases. Gray lines represent location of amplicons. **(b)** Sorted log10 coverage of each base. Black solid line represents the median coverage, green shade represents 0.2× to 5× median coverage (Olivar: 46.3% bases, ARTIC v4.1: 42.6% bases), olive shade represents 0.1× to 10× coverage (Olivar: 64.1% bases, ARTIC v4.1: 59.3% bases), red dashed line represents 0.05× median coverage (Olivar: 19.1% bases less than 0.05×, ARTIC v4.1: 17.3% bases less than 0.05×).

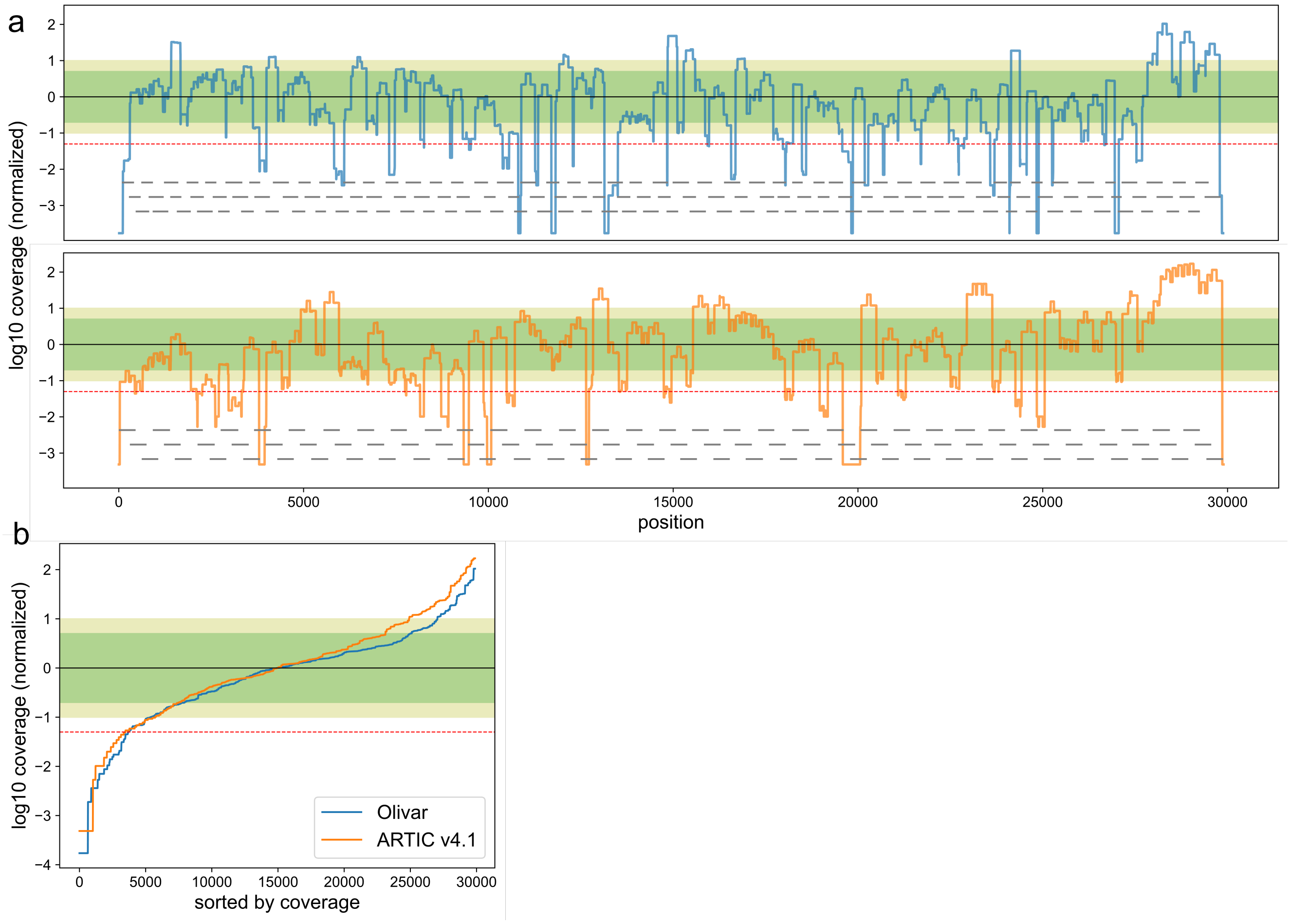


**Figure 8.** **Sars-Cov-2 whole genome coverage of both Olivar (blue) and ARTIC v4.1 (orange) primers. Figures showing results from one wastewater sample (site: KB, Aug. 15, 2022).** **(a)** log10 coverage of each base. Coverage is normalized by median coverage of all bases. Gray lines represent location of amplicons. **(b)** Sorted log10 coverage of each base. Black solid line represents the median coverage, green shade represents 0.2× to 5× median coverage (Olivar: 56.9% bases, ARTIC v4.1: 52.0% bases), olive shade represents 0.1× to 10× coverage (Olivar: 72.4% bases, ARTIC v4.1: 64.2% bases), red dashed line represents 0.05× median coverage (Olivar: 12.4% bases less than 0.05×, ARTIC v4.1: 11.6% bases less than 0.05×).
